# Decorating chromatin for enhanced genome editing using CRISPR-Cas9

**DOI:** 10.1101/2022.03.15.484540

**Authors:** Evelyn Chen, Enrique Lin-Shiao, Marena Trinidad, Mohammad Saffari Doost, David Colognori, Jennifer A. Doudna

**Affiliations:** Department of Molecular and Cell Biology, University of California, Berkeley, Berkeley, CA USA 94720; Innovative Genomics Institute, University of California, Berkeley, Berkeley, CA USA 94720; Howard Hughes Medical Institute, University of California, Berkeley, Berkeley, CA USA 94720; Department of Chemistry, University of California, Berkeley, Berkeley, CA USA 94720; Molecular Biophysics and Integrated Bioimaging Division, Lawrence Berkeley National Laboratory, Berkeley, CA USA 94720; California Institute for Quantitative Biosciences (QB3), University of California, Berkeley, Berkeley, CA USA 94720; Gladstone Institutes, University of California, San Francisco, CA USA 94158; Gladstone-UCSF Institute of Genomic Immunology, San Francisco, CA USA 94158

**Keywords:** CRISPR-Cas9, genome editing, chromatin, epigenetics

## Abstract

CRISPR-associated (Cas) enzymes have revolutionized biology by enabling RNA-guided genome editing. Homology-directed repair (HDR) in the presence of donor templates is currently the most versatile method to introduce precise edits following CRISPR-Cas-induced double-stranded DNA cuts, but HDR efficiency is generally low relative to end-joining pathways that lead to insertions and deletions (indels). We tested the hypothesis that HDR could be increased using a Cas9 construct fused to PRDM9, a chromatin remodeling factor that deposits histone methylations H3K36me3 and H3K4me3 to mediate homologous recombination in human cells. Our results show that the fusion protein contacts chromatin specifically at the Cas9 cut site in the genome to increase the observed HDR efficiency by three-fold and HDR:indel ratio by five-fold compared to that induced by unmodified Cas9. HDR enhancement occurred in multiple cell lines with no increase in off-target genome editing. These findings underscore the importance of chromatin features for the balance between DNA repair mechanisms during CRISPR-Cas genome editing and provide a new strategy to increase HDR efficiency.

**Significance Statement:** CRISPR-Cas-mediated homology-directed repair (HDR) enables precision genome editing for diverse research and clinical applications, but HDR efficiency is often low due to competing end-joining pathways. Here, we describe a simple strategy to influence DNA repair pathway choice and improve HDR efficiency by engineering CRISPR-Cas9-methyltransferase fusion proteins. This strategy highlights the impact of histone modifications on DNA repair following CRISPR-Cas-induced double-stranded breaks and adds to the CRISPR genome editing toolbox.

## Introduction

CRISPR-Cas9 genome editing provides a transformative opportunity to study and treat a wide range of diseases (1–4). CRISPR enzymes generate RNA-guided double-stranded breaks (DSBs) in DNA that initiate repair by mechanisms including non-homologous end joining (NHEJ), microhomology-mediated end joining (MMEJ), and homology-directed repair (HDR) (5). NHEJ and MMEJ pathways lead to a heterogeneous mixture of insertions and deletions (indels), which have been harnessed to facilitate gene disruption. HDR in the presence of donor templates can produce defined genomic changes, but its low efficiency relative to NHEJ and MMEJ repair remains a limitation of genome editing applications (6). Cas9-based methods including base editing (7, 8) and prime editing (9) enable targeted substitutions and small insertions without requiring a DSB. However, these approaches are limited to single nucleotide substitutions or insertions smaller than 50 bases and have also been associated with off-target RNA editing (10). Therefore, HDR continues to be the most versatile method for targeted substitutions and insertions.

Previous studies of the effects of chromatin on DNA repair outcomes following CRISPR-Cas9-induced DSBs have focused primarily on the differences between euchromatin (accessible chromatin) and heterochromatin (tightly packaged and inaccessible chromatin) (11, 12). Heterochromatin temporarily inhibits targeting by CRISPR-Cas9, but the impact of chromatin context on DNA repair pathway choice following the initial delay remains poorly understood (13–15). It has been suggested that NHEJ frequencies following CRISPR-Cas9-induced DSBs are higher at euchromatic regions whereas MMEJ and HDR are biased toward heterochromatin (16, 17), although one study found no significant influence of preexisting chromatin contexts on the relative frequencies of DNA repair pathways (15).

The balance between HDR and end-joining pathways has been correlated with locus-specific chromatin features. H3K36 trimethylation (H3K36me3) is a histone modification critical for homologous recombination (HR) in human cells (18–20). Previous studies have shown that genomic sites marked with H3K36me3 are preferentially repaired by HR following AsisI-induced DSBs (20, 21). However, the role of site-specific H3K36me3 in DNA repair of CRISPR-Cas9-induced DSBs has remained largely unexplored.

In this study, we investigated how pre-existing and newly deposited H3K36me3 affect the efficiency of CRISPR-Cas9-mediated HDR by engineering and testing fusion proteins of Cas9 to H3K36 methyltransferases involved in DSB repair. We selected PRDM9 as it deposits both H3K36me3 and H3K4me3 during meiotic recombination to regulate DSB hotspot localization and repair (Fig. S1A) (22, 23). During meiosis, SPO11 is recruited to PRDM9-binding sites to catalyze DSB formation, and DMC1 mediates homologous strand invasion and crossovers (22, 23). ZCWPW1, an epigenetic reader recently found to have co-evolved with PRDM9, specifically recognizes the dual histone methylation marks and is required for synapsis and DSB repair (Fig. S1A) (24–27). Furthermore, PRDM9 is expressed exclusively in germ cells during meiosis, and previous work in mice demonstrated that it does not influence transcription, making it an attractive fusion partner for Cas9 editing tool development (28). We also tested a fusion of Cas9 to SETD2, which deposits H3K36me3 in somatic cells to recruit CtIP to DSB sites, promoting DNA end resection and RPA and RAD51 foci formation which stimulates accurate genome repair (Fig. S1A) (19). SETD2-mediated H3K36me3 deposition also enables V(D)J recombination in the adaptive immune system and broadly decorates transcriptionally active regions to ensure transcriptional fidelity (Fig. S1A) (19, 29). Lastly, we fused Cas9 to SETMAR (Metnase), a H3K36me2 methyltransferase that has been suggested to mediate NHEJ and suppress chromosomal translocations (Fig. S1A) (30–32).

We show that both endogenous and newly deposited histone modifications influence DNA repair pathway choice. In particular, the presence of H3K36me3 favors DSB repair via HDR following CRISPR-Cas9 cuts. We explored the efficacy of PRDM9-Cas9-catalyzed genome editing across different cell types, demonstrating a three-fold increase in HDR rate and a five-fold increase in the HDR:indel ratio relative to that observed using unmodified Cas9.

These findings underscore the importance of chromatin modifications for DNA repair pathway choice during CRISPR-Cas9-mediated genome editing and provide a new approach to enhance HDR efficiency.

## Results

### Endogenous chromatin architecture modulates DNA repair pathway choice

To assess whether endogenous chromatin architecture affects Cas9 activity and DNA repair pathway choice, we examined H3K36me3 and H3K4me3 profiles based on publicly available ENCODE ChIP-seq data from human embryonic kidney 293 (HEK293) cells (33). We selected 16 target sites with varying H3K36me3 and H3K4me3 enrichment, including a disease-relevant site C9 in *LDLR* which encodes the low-density lipoprotein receptor (Fig. 1A). We measured HDR frequencies at each site by transfecting HEK293T cells with Cas9 and single guide RNA (sgRNA) expression plasmids along with a single-stranded oligodeoxynucleotide (ssODN) donor template encoding a point mutation upstream of the protospacer adjacent motif (PAM), a short DNA sequence adjacent to the sgRNA binding site at each locus (34). At 3 days post-transfection (dpt), we evaluated the extent of genome modification by next-generation sequencing (NGS).

**Fig. 1:**
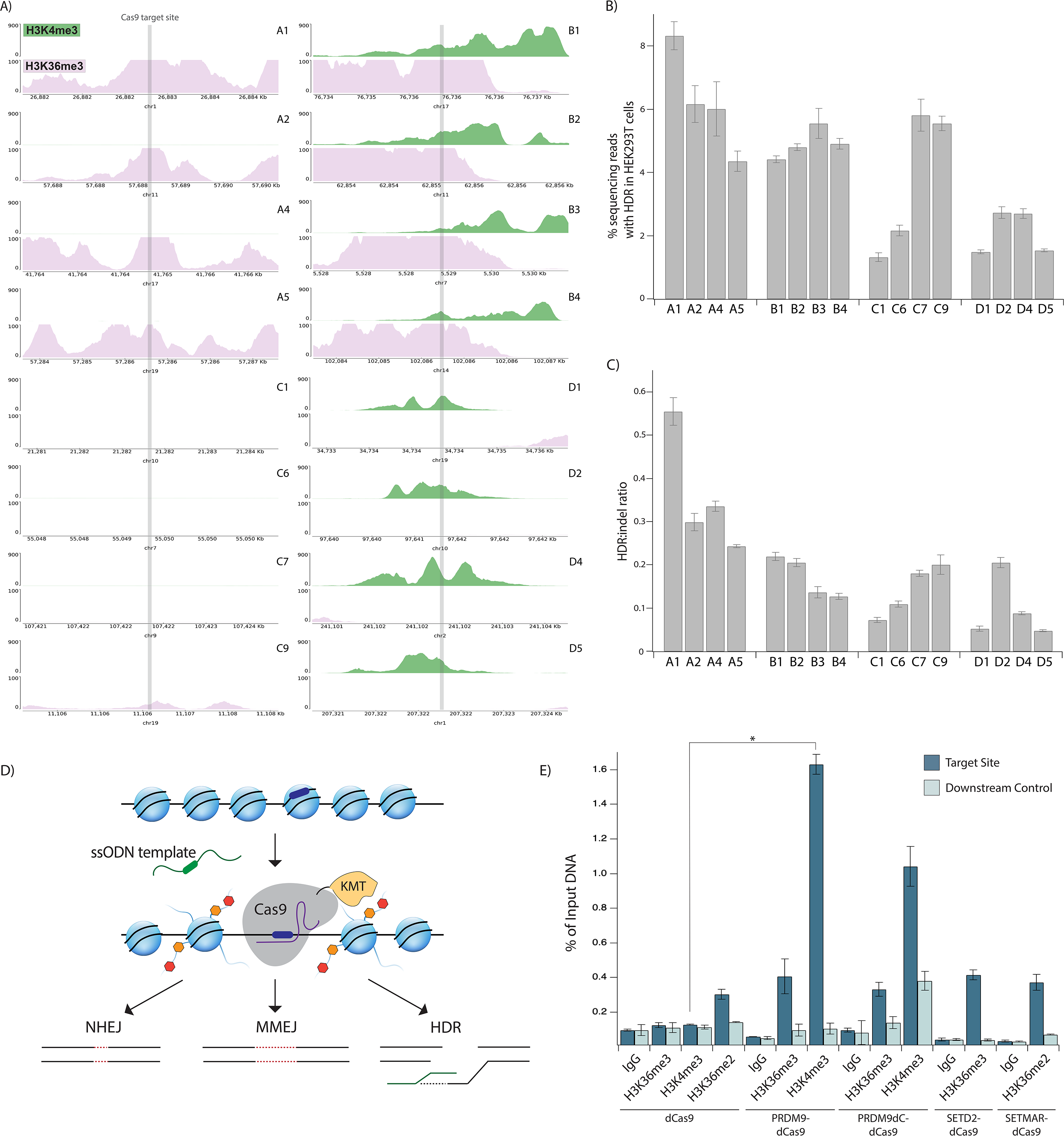
Endogenous histone modifications mediate DNA repair pathway choice. A) UCSC genome browser tracks showing normalized endogenous levels of H3K4me3 (green) and H3K36me3 (pink) at target site ± 1.5 kilobases (kb) based on ENCODE ChIP-seq datasets from HEK293 cells. Each plot spans 3□kb and the y-axis reports the negative-log p-value for peak enrichment. B) HDR frequency measured by next-generation sequencing (NGS) at genomic sites with varying endogenous H3K4me3 and H3K36me3 enrichment in HEK293T cells transfected with plasmids expressing Cas9 and sgRNA along with a ssODN template with 50-nt homology arms. Data represents mean ± standard deviation (s.d.) (n = 3). C) HDR:indel ratio measured by NGS at genomic sites with varying endogenous H3K4me3 and H3K36me3 enrichment in HEK293T cells transfected with plasmids expressing Cas9 and sgRNA along with a ssODN template with 50-nt homology arms. Data represents mean ± s.d. (n = 3). D) Schematic of Cas9-histone lysine methyltransferase (KMT) fusion protein activity. Cas9 fusion protein is guided by a sgRNA (purple) to the DNA target site (dark blue), which may be embedded within a heterochromatic region. Cas9 fusion protein deposits histone marks (orange and red) at the target site, which influence the choice of DNA repair pathway following Cas9-induced DSBs. E) H3K36me3, H3K4me3, and H3K36me2 enrichment shown as a percentage of input DNA measured by ChIP-qPCR at site C7 in HEK293T cells at 3 days post-transfection (dpt). Cells were transfected with plasmids expressing the dCas9 fusion proteins and sgRNA. Data represents mean ± s.d. (n = 2). \**P* < 0.05, determined by student’s two-tailed t-test.

We observed that while HDR frequencies are not strongly correlated with H3K36me3 or H3K4me3 enrichment, sites decorated with either H3K36me3 only (sites A1, A2, A4, A5) or both H3K36me3 and H3K4me3 (sites B1-B4) showed higher HDR rates on average than those at sites lacking H3K36me3 or H3K4me3 (sites C1, C6, C7, C9) (Fig. 1B). Of note, site A1, which is extensively marked with H3K36me3, showed the highest HDR efficiency (8.3 ± 0.6%) among all target sites, whereas site C1, which lacks either histone modifications, had the lowest HDR efficiency (1.3 ± 0.1%) (Fig. 1B). Similarly, HDR:indel ratios were highest at sites enriched with H3K36me3 only, with site A1 showing a HDR:indel ratio of 0.55 ± 0.03, and unenriched sites showed lower HDR:indel ratios on average in comparison (Fig. 1C). Interestingly, sites enriched with H3K4me3 only (sites D1, D2, D4, D5) displayed the lowest HDR efficiencies and HDR:indel ratios on average (Fig. 1B-C). Given previous studies showing that H3K4me3 decreases upon DSB induction via AsiSI to facilitate HR repair (35, 36), these findings suggest that H3K4me3 may also hinder HDR following Cas9-induced cuts. Together, our results indicate that endogenous H3K36me3 favors HDR over error-prone end-joining pathways following Cas9-induced DSBs compared to sites without H3K36me3.

### PRDM9-Cas9 fusion proteins modify histones site-specifically

Based on the above findings and previous reports (14, 37), we hypothesized that newly deposited histone modifications might influence DNA repair pathway choice following Cas9-induced DNA cutting (Fig. 1D). To test this idea, we constructed genes encoding four chimeric proteins in which histone methyltransferases are fused at the N-terminus of Cas9 in the pCAGGS expression vector (Fig. S1B). The PRDM9-Cas9 fusion comprises the KRAB domain which recruits additional proteins to facilitate recombination, the PR/SET domain which catalyzes methyltransferase activity, and a post-SET single zinc finger (ZnF) (28, 38). The N-terminal domains of PRDM9 may be important for mediating interactions with chromatin remodeling factor HELLS and forming a pioneer complex to open chromatin (39). Based on RNA sequencing (RNA-seq) analysis, we determined that HELLS is expressed in HEK293T cells (Fig. S1C). We excluded the C-terminal ZnF array of PRDM9 to avoid its endogenous DNA binding activity (40). The post-SET ZnF of PRDM9 has been proposed to be involved in the autoregulation of methyltransferase activity (38), although a previous study found that PRDM9 without the post-SET ZnF did not lead to higher methylation activity (28). To further investigate this, we also engineered PRDM9dC-Cas9, a truncated version lacking the post-SET ZnF (Fig. S1B). RNA-seq analysis showed that ZCWPW1, which recognizes PRDM9-mediated histone methylation marks to mediate successful DSB repair during meiosis, is not highly expressed in HEK293T cells (Fig. S1C) (24–27). SETD2-Cas9 includes the SET and post-SET domains of SETD2 which deposit H3K36me3 (Fig. S1B) (19). SETMAR-Cas9 includes the SET domain of SETMAR, important for NHEJ repair (Fig. S1B) (30, 31).

We performed Western blot analysis to determine the level of expression of each fusion protein compared to unmodified Cas9 in HEK293T cells. When we delivered equal amounts (500ng) of plasmids expressing either Cas9 or the fusion proteins, we observed slightly lower expression of each fusion compared to Cas9 at 3 dpt (Fig. S1D). Notably, scaling the amounts of plasmids expressing the fusion proteins based on their size compared to unmodified Cas9 did not lead to increased expression (Fig. S1D). We reasoned that the lower expression of fusion constructs could be due to the larger protein sizes or decreased stability.

We then investigated whether the Cas9 fusion proteins deposit histone modifications in a site-specific manner using chromatin immunoprecipitation followed by quantitative PCR (ChIP-qPCR). To eliminate the potential effects of DSB remodeling on chromatin modifications, we engineered fusion proteins of nuclease-dead Cas9 (dCas9) to each histone methyltransferase described above. We transfected HEK293T cells with either dCas9 or a dCas9 fusion protein and sgRNAs targeting site C7 or C9 and analyzed H3K36me3, H3K4me3, and H3K36me2 signals at 3 dpt.

Strikingly, at site C7, PRDM9-dCas9 and PRDM9dC-dCas9 increased H3K36me3 by up to 3.2-fold and H3K4me3 by up to 12.7-fold compared to dCas9 (Fig. 1E). The downstream region of site 7 did not show increased histone modifications by PRDM9-dCas9 compared to dCas9, indicating that the fusion protein specifically deposited histone marks at the Cas9-targeted site (Fig. 1E). Although PRDM9dC-dCas9 slightly increased H3K36me3 and H3K4me3 at the downstream control site, the increase was not statistically significant (Fig. 1E). In addition, SETD2-dCas9 increased H3K36me3 by 3.3-fold, and SETMAR-dCas9 increased H3K36me2 by 1.2-fold compared to dCas9 at site C7 without modifying the downstream control site (Fig. 1E).

We observed preexisting H3K36me3 at site C9, consistent with ENCODE ChIP-seq data from HEK293 cells (Fig. S1E). PRDM9 fusion proteins deposited H3K36me3 marks at site 9 (1.5-fold enrichment by PRDM9-dCas9 and 1.7-fold enrichment by PRDM9dC-dCas9), and they increased H3K4me3 by up to 17.1-fold (Fig. S1E). SETD2-dCas9 increased H3K36me3 by 4.2-fold, and SETMAR-dCas9 increased H3K36me2 by 1.4-fold (Fig. S1E). The modest increase in H3K36me3 at site C9 by the fusion proteins may be specific to this locus given its preexisting H3K36me3. In addition, the low methylation activity of SETMAR-dCas9 at site C7 and C9 may be due to preexisting H3K36me2 at both targets. Overall, PRDM9-dCas9 deposited histone modifications at target sites with the highest efficiency and specificity among the fusion protein designs.

### CRISPR-Cas9-methyltransferase fusion proteins display higher HDR and HDR:indel ratios

We next evaluated the editing outcomes of the fusion constructs via a BFP-to-GFP conversion assay in HEK293T cells stably expressing a BFP reporter (41). Cells were transfected with plasmids encoding either Cas9 or one of the four fusion proteins and a sgRNA targeting the BFP gene, as well as a ssODN template encoding a three-nucleotide (nt) change. Cells express GFP if the Cas9-induced DSB is repaired via HDR; on the other hand, cells lose BFP expression if the cut is repaired via end-joining pathways leading to indel formation (Fig. 2A). BFP-/GFP+ (HDR) and BFP-/GFP- (indel) cells were gated relative to cells transfected with a non-targeting control (NTC) sgRNA (Fig. 2B).

**Fig. 2:**
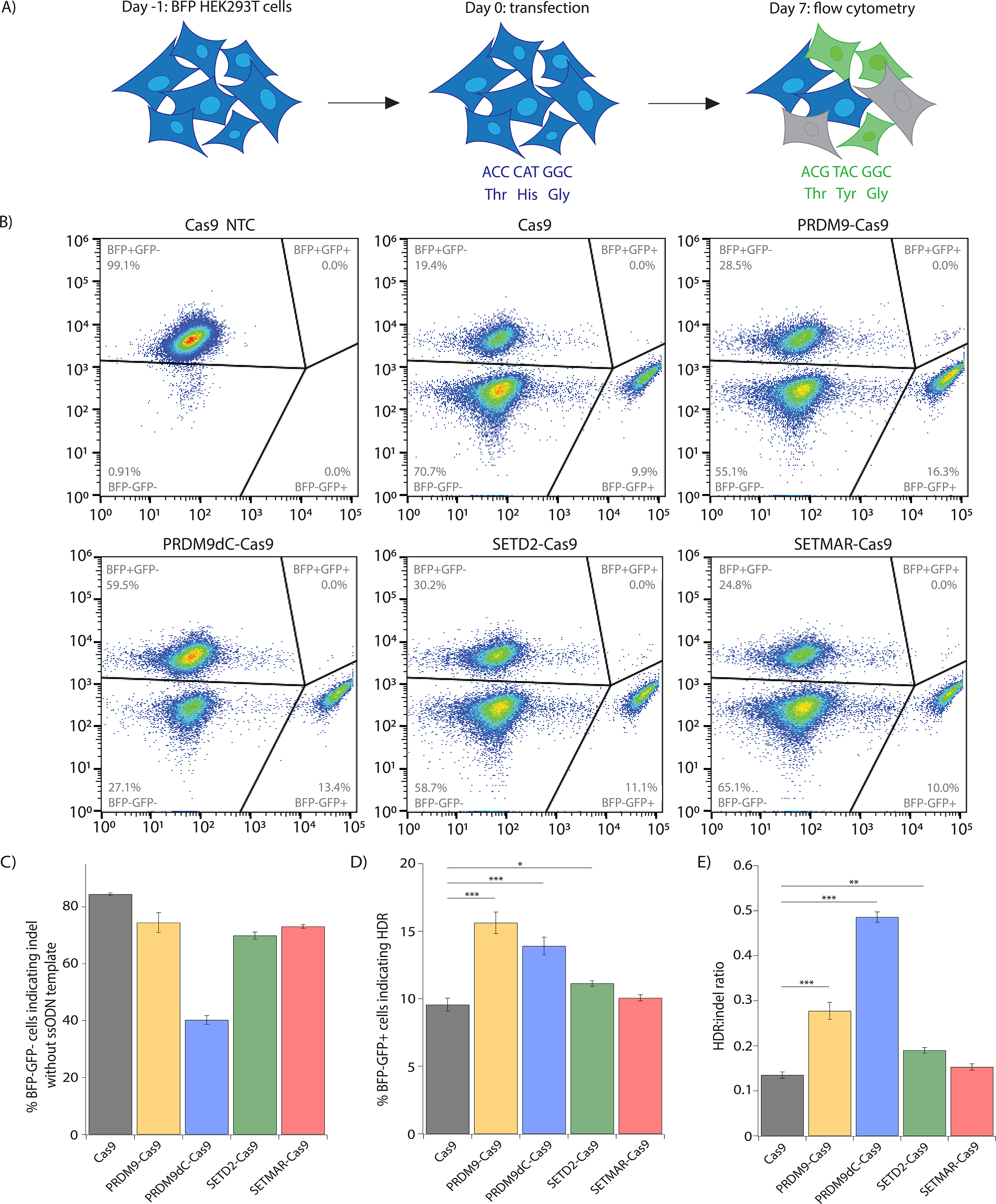
Engineered CRISPR-Cas9-methyltransferase fusion proteins produce higher HDR and HDR:indel ratios relative to Cas9. A) Schematic of BFP-to-GFP reporter assay. In brief, BFP+/GFP- HEK293T cells can be converted to BFP-/GFP+ cells via HDR or to BFP-/GFP- via indel formation. Flow cytometry was performed 7 days post-transfection (dpt). B) Flow cytometry plots for BFP-to-GFP reporter cells showing frequency of HDR (BFP-/GFP+) and indel (BFP-/GFP-). Cells were transfected with plasmids expressing the fusion proteins and sgRNA along with a ssODN template with a 91nt homology arm on the PAM-proximal side of the DSB and a 36nt homology arm on the PAM-distal side. A non-targeting negative control (NTC) sgRNA was included. Representative plots are shown for 1 biological replicate. C) Editing activity of Cas9-methyltransferase fusion proteins measured by flow cytometry in BFP-to-GFP reporter cells transfected with plasmids encoding fusion protein and sgRNA, without ssODN template. D) HDR frequency measured by flow cytometry in BFP-to-GFP reporter cells transfected with plasmids encoding fusion protein and sgRNA, along with ssODN template. E) HDR:indel ratio measured by flow cytometry in BFP-to-GFP reporter cells transfected with plasmids encoding fusion protein and sgRNA, along with ssODN template. For C-E, data represents mean ± s.d. (n = 4). \**P* < 0.05, \*\**P* < 0.01, \*\*\**P* < 0.001, determined by student’s two-tailed t-test.

Using flow cytometry at 7 dpt, we observed that in the absence of HDR templates, the Cas9 fusion proteins induced modestly reduced genome edits relative to those induced by unmodified Cas9, except for PRDM9dC-Cas9 which displayed a 50% decrease in editing efficiency (Fig. 2C). This could result from reduced nuclease activity of the Cas9 fusion proteins or increased deposition of histone modifications that improved the rate of perfect DNA repair following Cas9-catalyzed cutting. Strikingly, when ssODN templates were co-introduced, PRDM9-Cas9 and PRDM9dC-Cas9 significantly increased HDR efficiency (15.6 ± 0.8% and 13.9 ± 0.7%) compared to unmodified Cas9 (9.6 ± 0.5%) (Fig. 2D). Additionally, PRDM9-Cas9 displayed a two-fold higher HDR:indel ratio (0.28 ± 0.02) compared to Cas9 (0.14 ± 0.01), and PRDM9dC-Cas9 achieved a nearly four-fold higher HDR:indel ratio (0.48 ± 0.01) (Fig. 2E). SETD2-Cas9 also increased HDR efficiency (11.1 ± 0.2%) and HDR:indel ratio (0.19 ± 0.01) compared to Cas9. Lastly, SETMAR-Cas9 did not modify the balance between HDR and end-joining pathways, possibly due to its low methylation activity. Overall, the PRDM9 fusion proteins significantly improved HDR-mediated DNA repair while decreasing indel formation.

### PRDM9-Cas9 fusion protein enhances HDR:indel ratios at endogenous genomic loci and in different cell types

We examined the impact of the Cas9 fusion proteins on DSB repair pathways at endogenous genomic sites. To this end, we transfected HEK293T cells with fusion protein and sgRNA expression plasmids and ssODN templates and measured the extent of genome modification by NGS. PRDM9-Cas9, PRDM9dC-Cas9, and SETD2-Cas9 all produced higher HDR levels than unmodified Cas9 at both site C7 and C9 (Fig. 3A-B). Notably, PRDM9-Cas9 displayed HDR frequencies of 11.0 ± 0.2% and 9.2 ± 0.6% at sites C7 and C9, compared to 5.8 ± 0.5% and 5.6 ± 0.2% with Cas9 at the respective sites (Fig. 3A-B). Moreover, PRDM9-Cas9 and SETD2-Cas9 improved HDR:indel ratios by three-fold or more compared to unmodified Cas9 (Fig. 3A-B).

**Fig. 3:**
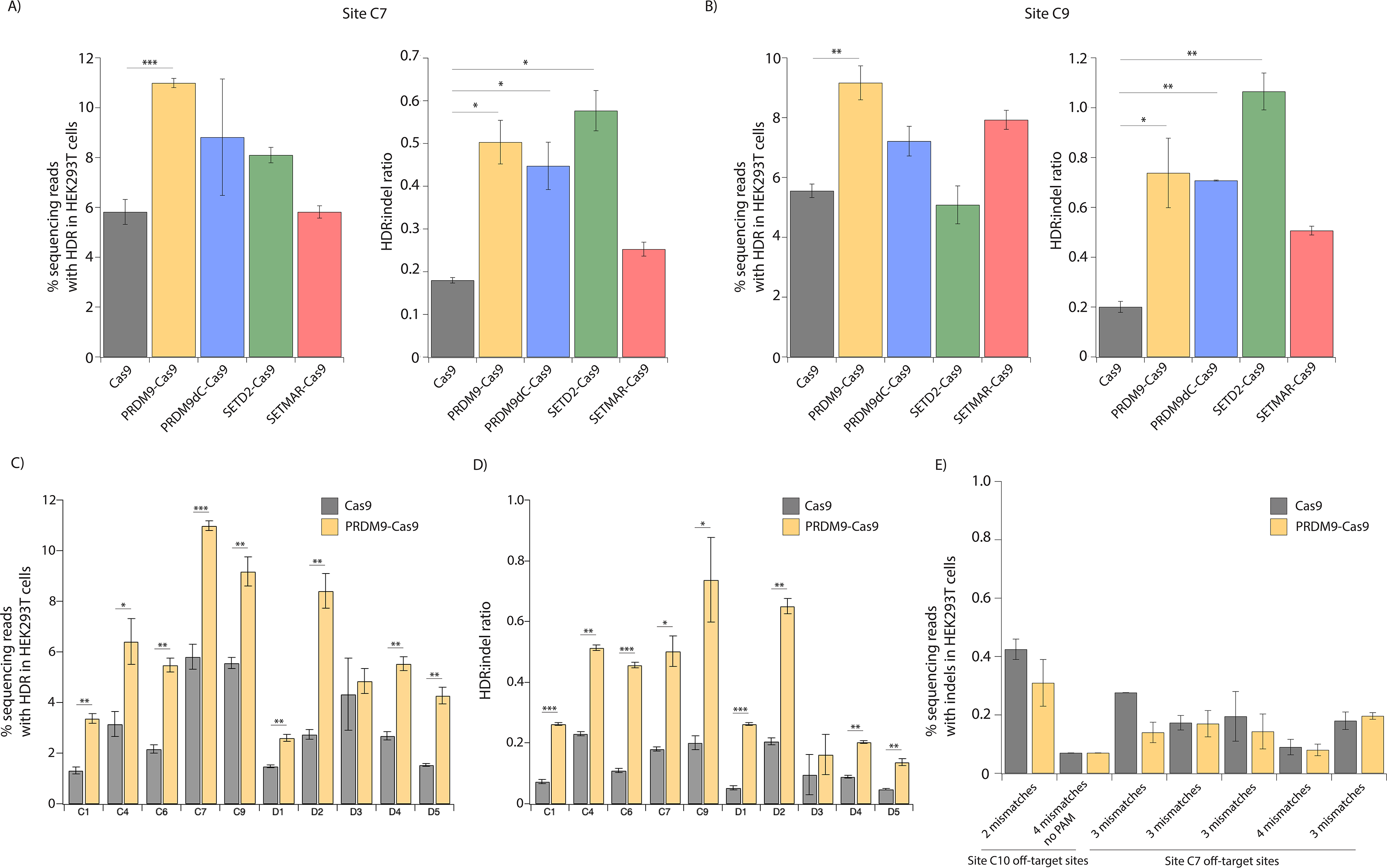
PRDM9-Cas9 fusion protein displays increased HDR efficiency and HDR:indel ratio across multiple genomic sites. A) HDR frequency and HDR:indel ratio measured by NGS at site C7 (intergenic) in HEK293T cells transfected with plasmids expressing fusion protein and sgRNA along with a ssODN template with 50-nt homology arms. B) HDR frequency and HDR:indel ratio measured by NGS at site C9 (exon 5 of *LDLR*) in HEK293T cells transfected with fusion protein and sgRNA along with a ssODN template with 50-nt homology arms. C) HDR frequency measured by NGS at ten genomic loci in HEK293T cells transfected with PRDM9-Cas9 and sgRNA with ssODN template. D) HDR:indel ratio measured by NGS at ten genomic loci in HEK293T cells transfected with PRDM9-Cas9 and sgRNA with ssODN template. E) Off-target activity of PRDM9-Cas9 measured by NGS at 7 potential off-target sites predicted from 2 sgRNAs. For A-D, data represents mean ± s.d. (n = 3). \**P* < 0.05, \*\**P* < 0.01, \*\*\**P* < 0.001, determined by student’s two-tailed t-test.

Since PRDM9-Cas9 generated the highest HDR frequency at both sites C7 and C9, we tested this fusion construct at additional genomic loci without H3K36me3 and H3K4me3 or with H3K4me3 only, where unmodified Cas9 achieved low HDR frequencies and HDR:indel ratios. PRDM9-Cas9-induced HDR frequencies were up to 2.5-fold higher than those induced by Cas9 at sites C1, C4, and C6 (Fig. 3C). Interestingly, PRDM9-Cas9 resulted in the largest increases in HDR frequency at H3K4me3-rich sites (D1-D5), achieving 8.4 ± 0.6% at site D2, which was 3.1-fold higher compared to Cas9 (Fig. 3C). PRDM9-Cas9 consistently improved HDR:indel ratios, resulting in a 4.2-fold increase in HDR:indel ratio at site C6 (0.46 ± 0.01 by PRDM9-Cas9 and 0.11 ± 0.01 by Cas9) and a 5.0-fold increase at site D1 (0.26 ± 0.003 by PRDM9-Cas9 and 0.05 ± 0.01 by Cas9) (Fig. 3D). At H3K36me3-rich sites (A1-A5), we detected up to 1.6-fold higher HDR levels when using PRDM9-Cas9 compared to Cas9 (Fig. S2A). PRDM9-Cas9 also showed higher HDR frequencies than Cas9 at four of the five sites enriched with both H3K36me3 and H3K4me3 (B1-B5) (Fig. S2C). HDR:indel ratios were significantly higher (1.4-fold to 3.1-fold increase) using PRDM9-Cas9 at four of the five sites with high H3K36me3 and all of the sites with both H3K36me3 and H3K4me3 (Fig. S2B,D). These findings establish that PRDM9-Cas9 improves HDR efficiency and HDR:indel ratio following Cas9-induced DSBs, especially at sites lacking H3K36me3 and H3K4me3 and sites marked with H3K4me3.

We then selected additional genomic sites without H3K36me3 or H3K4me3 enrichment, including a disease-relevant site (C8) in the *HBB* gene which encodes the beta-globin component of hemoglobin, to test PRDM9-Cas9 (Fig. S3). Unexpectedly, the fusion protein did not improve HDR efficiency or HDR:indel ratio at sites C10 and C11 (Fig. S2E-F). Since the baseline HDR frequencies at C10 and C11 achieved by unmodified Cas9 were very high, it is possible that HDR cannot be further improved by depositing H3K36me3. Nonetheless, PRDM9-Cas9 achieved higher HDR frequencies at five of the eight additional sites compared to Cas9.

To investigate the limited effect of PRDM9-Cas9 at some target sites, we considered that other chromatin features may impact the relative frequencies of HDR and end-joining pathways. Based on the availability of ENCODE ChIP-seq datasets, we examined endogenous levels of H3K79me2 and H4K20me1 which are more abundant at baseline near AsisI-induced DSBs preferentially repaired via HR (21), and H3K9me3 which is associated with heterochromatin (Fig. S4A-C) (33, 42). Sites marked with H3K36me3 and H3K4me3 (B1-B5) also showed H3K79me2 and H4K20me1 enrichment, and sites A3 and A5 and several sites without H3K36me3 or H3K4me3 showed moderate levels of H3K9me3 (Fig. S4A-B). We observed that Cas9-induced HDR frequency and HDR:indel ratio were not strongly associated with these additional histone modifications. Sites C10 and C11 were not marked with H3K79me2 or H4K20me1, suggesting that other factors, such as additional chromatin features and sgRNA design, may be favoring HDR at these sites (Fig. S4B-C) (21, 43).

Given that the Cas9 fusion constructs can modify adjacent chromatin, we then examined whether this could lead to higher off-target editing by evaluating editing activity at seven potential off-target sites associated with two well-characterized sgRNA sequences (44). Importantly, PRDM9-Cas9 does not lead to higher off-target editing compared to Cas9 (Fig. 3E). Together, these data highlight the ability of PRDM9-Cas9 to improve Cas9-mediated HDR efficiency and HDR:indel ratio via *de novo* modifications of chromatin architecture without increasing off-target effects.

Next, we compared PRDM9-Cas9- and Cas9-induced HDR efficiency using ssODN templates that either include or lack a mutation at the PAM site. Previous reports have shown that ssODNs that introduce a blocking mutation at the PAM significantly increase HDR efficiency by preventing the retargeting of the edited site (45, 46). We therefore tested ssODN templates that either disrupt or retain the PAM sequence at sites C7, C9, and C10. Consistent with previous reports, unmodified Cas9 showed two-fold higher HDR efficiency at sites C7 and C9 using ssODNs with PAM mutations (10.6% ± 0.3% and 12.3% ± 0.5%) compared to ssODNs without PAM mutations (5.8% ± 0.5% and 5.6% ± 0.2%) although no significant difference was observed at site C10 (Fig. 4A) (45, 46). Of note, our PRDM9-Cas9 fusion approach alone achieved similar improvements in HDR efficiency compared to unmodified Cas9 using ssODN templates with PAM mutations (Fig. 4A). PRDM9-Cas9 increased the HDR:indel ratio more significantly than the PAM mutation strategy (Fig. 4B). Taken together, our findings suggest that PRDM9-Cas9 can be utilized to improve HDR efficiency effectively without the need to introduce PAM mutations.

**Fig. 4.**
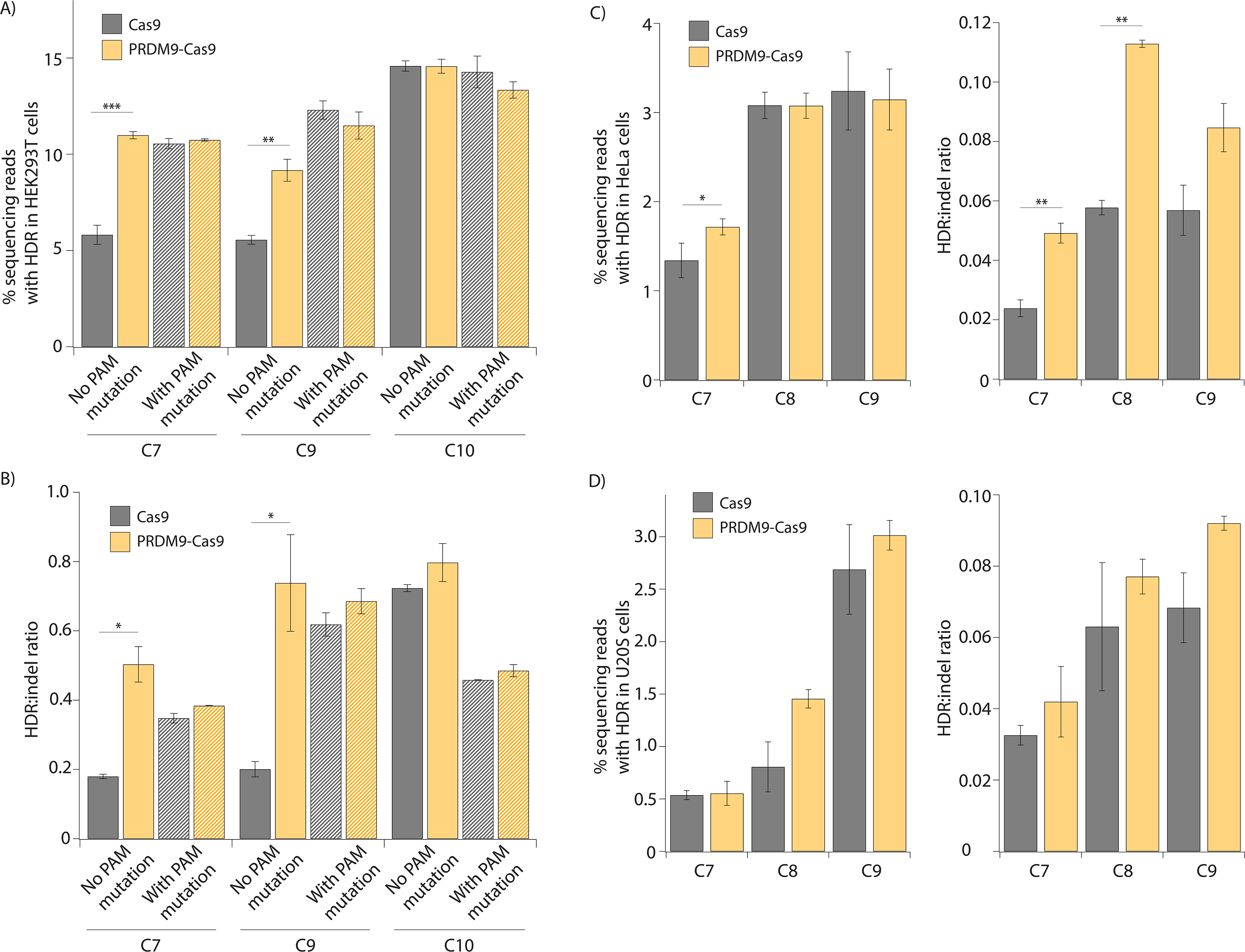
PRDM9-Cas9 fusion protein produces increased HDR:indel ratios in different cell types. A) HDR frequency measured by NGS at multiple genomic loci (C7, C9, C10) in HEK293T cells transfected with PRDM9-Cas9 and sgRNA with ssODN templates that either include (hash pattern) or lack (solid pattern) a mutation at the PAM site. B) HDR:indel ratio measured by NGS at multiple genomic loci (C7, C9, C10) in HEK293T cells transfected with PRDM9-Cas9 and sgRNA with ssODN templates that either include (hash pattern) or lack (solid pattern) a mutation at the PAM site. C) HDR frequency and HDR:indel ratio measured by NGS at multiple genomic loci (C7-C9) in HeLa cells transfected with PRDM9-Cas9 and sgRNA with ssODN template. D) HDR frequency and HDR:indel ratio measured by NGS at multiple genomic loci (C7-C9) in U2OS cells transfected with PRDM9-Cas9 and sgRNA with ssODN template. For A-D, data represents mean ± s.d. (n = 3). \**P* < 0.05, \*\**P* < 0.01, \*\*\**P* < 0.001, determined by student’s two-tailed t-test.

To investigate whether PRDM9-Cas9 can improve HDR efficiency in other cell types, we tested the fusion construct at sites C7-C9 in HeLa and U2OS cells. We only observed a modest increase in HDR frequency by PRDM9-Cas9 in these cell types, which may be due to differences in histone modifications and DNA repair processes that are cell type-specific (Fig. 4C-D). Nonetheless, PRDM9-Cas9 enhanced HDR:indel ratios by up to two-fold across all three target sites in both HeLa and U2OS cells (Fig. 4C-D). Overall, PRDM9-Cas9 improved the frequency of HDR relative to indels at multiple endogenous sites across different mammalian cell lines.

## Discussion

HDR-mediated precision genome editing holds great potential for research and clinical applications, but HDR efficiency is often low and considerably variable across different genomic loci and cell types. Here, we report a simple strategy to influence DNA repair pathway choice and improve HDR efficiency by engineering CRISPR-Cas9-methyltransferase fusion proteins to deposit histone modifications involved in HR during meiosis and somatic DNA repair (18–23). Among the four fusion constructs tested, PRDM9-Cas9 deposited H3K36me3 and H3K4me3 specifically at target sites and improved HDR efficiency by up to three-fold and HDR:indel ratio by up to five-fold at multiple genomic loci without increasing off-targeting editing. The consistent improvements in HDR:indel ratio achieved by PRDM9-Cas9 makes it particularly valuable for precise genome editing applications that demand a high HDR frequency relative to indel formation, such as gene correction in sickle cell disease (47). The lower observed indel formation may be explained by increased DSB repair via sister chromatid or interhomolog homologous recombination due to de novo H3K36me3. Additionally, it was previously determined that ectopic PRDM9 expression in HEK293 cells can lead to the recruitment of HELLS to PRDM9-binding sites (39). Therefore, our PRDM9-Cas9 fusion construct, which includes the N-terminal domains of PRDM9, may also interact with HELLS to mediate chromatin remodeling at Cas9 target sites, potentially influencing the choice of DSB repair pathway in the present study. Given these possibilities, further studies are needed to characterize the mechanism in which PRDM9-Cas9 increases HDR efficiency and HDR:indel ratio.

While PRDM9-Cas9 achieved a clear improvement in HDR efficiency at most target sites lacking endogenous H3K36me3 enrichment, the effect was not observed across all sites investigated. This could be explained by other site-specific histone modifications and factors beyond chromatin architecture that affect DSB repair at different loci and in different cell types. In particular, a previous study reported that genomic sites endogenously enriched for H4K16 acetylation (H4K16ac) are associated with increased HDR of I-SceI-induced DNA cuts (48). Other pre-existing histone modifications associated with DSBs preferentially repaired by HDR include H3K79me2, H2BK120ub, H3K4me2, and H4K20me1 (21, 49). Accordingly, it is likely that the overall endogenous chromatin environment at certain target sites favors DSB repair via HDR such that the ability of PRDM9-Cas9 to improve HDR efficiency and HDR:indel ratio is limited beyond the levels achieved by unmodified Cas9.

It has been reported that PRDM9-mediated H3K36me3 exhibits lower enrichment and correlates negatively with H3K4me3 levels at highly active promoters (50). This property may impact the activity of PRDM9-Cas9 at target sites within promoters, warranting further work to investigate this possibility. However, since disease-associated single-nucleotide polymorphisms (SNPs) are more abundant in enhancers and other noncoding regions than promoters, PRDM9-Cas9 would be useful for correcting most clinically relevant SNPs at improved efficiency as demonstrated here (51).

Previous work has demonstrated that PRDM9 is not involved in transcriptional regulation *in vivo*, suggesting that PRDM9-Cas9 may not influence transcription while improving HDR (28). However, given the diversity of PRDM9 allelic variants (39, 52), additional studies are needed to evaluate whether our PRDM9-Cas9 fusion construct affects transcription. Nonetheless, PRDM9-Cas9 is not expected to result in persistent changes to the epigenome as studies have shown that CRISPR-based epigenome editing cannot be stably maintained without constitutive transgene expression or additional changes to DNA methylation status (53–55).

Donor templates that introduce additional point mutations to block the PAM site from re-cutting have been commonly used as an effective strategy to improve HDR efficiency (46). In this study, we directly compared the PRDM9-Cas9 fusion protein with the PAM-blocking strategy and observed similar increases in HDR efficiency relative to Cas9 alone. The fusion strategy has a significant advantage over PAM mutations as it enables the scarless introduction of a desired mutation at improved efficiency at a broader range of targets whereas a silent PAM-blocking mutation may not be available at some exonic target sites (46). Furthermore, mutating the PAM sequence in non-coding regions may disrupt transcription factor binding and downstream modulation of expression. While we did not observe a further increase in HDR when combining PRDM9-Cas9 with PAM-blocking mutations compared to using either strategy alone, this may be specific to the sites investigated. We anticipate that given the flexibility of fusion proteins, our fusion strategy can be combined with other newly developed strategies to potentially improve HDR efficiency. Importantly, recent studies reported on the impact of sgRNA design and CRISPR-Cas cut (blunt vs. staggered) on subsequent DNA repair pathway choice (43). Additionally, asymmetric donor templates and tiling sgRNAs have successfully improved HDR efficiency in several studies (41, 56).

Overall, our findings suggest that both endogenous and newly deposited histone modifications influence DNA repair pathway choice during CRISPR-Cas9-mediated genome editing. The engineered fusion proteins described here provide a transient strategy to enhance HDR efficiency by decorating adjacent nucleosomes with H3K36me3 and H3K4me3, allowing the development of better editing tools for research and therapeutic approaches.

## Materials and Methods

### Dataset

ChIP-seq datasets for H3K36me3 and H3K4me3 in HEK293 cells were obtained from the ENCODE portal with the following identifier: ENCSR372WXC (https://www.encodeproject.org/reference-epigenomes/ENCSR372WXC/). We selected target sites based on p-values for H3K4me3 and H3K36me3 enrichment. To better understand the chromatin landscape at each target site, we downloaded additional datasets from ENCODE (ENCFF315TAU, ENCFF714CDE, ENCFF758LNF, ENCFF127XFD, ENCFF502EIH) and observed the p-values for H3K9me3, H3K27me2 and H4K20me1 enrichment at each region.

### Plasmid construction

The Cas9 expression plasmid pCAGGS was a gift from Jennifer R. Hamilton. Cas9 was amplified from the expression plasmid using CloneAmp HiFi PCR Premix (Takara Bio) for 35 cycles (98□ for 10 s, 55□ for 15 s, and 72□ for 10 s; then 72□ for 1 min). PRDM9 (aa 1-416) and PRDM9dC (aa 1-371) were amplified from human cDNA purchased from GenScript (ORF Clone ID OHu03253). SETD2 (aa 1450-1645) was amplified from SETD2-GFP (Addgene plasmid # 80653; http://n2t.net/addgene:80653), and SETMAR (aa 14-277) was amplified from SETMAR (3BO5) (Addgene plasmid # 25250; http://n2t.net/addgene:25250). In-Fusion Cloning (Takara Bio) was used to clone PRDM9-Cas9, PRDM9dC-Cas9, SETD2-Cas9, and SETMAR-Cas9 into the pCAGGS expression vector. All Cas9 variants were confirmed by Sanger sequencing and are available on Addgene.

DNA oligonucleotides (IDT) for Cas9 sgRNA, including a non-targeting negative control sgRNA, were cloned into U6-sgRNA expression vectors (Table S1). Oligos were resuspended in nuclease-free water, and forward and reverse oligos were mixed with T4 polynucleotide kinase and 10X T4 DNA ligase buffer and placed in a thermal cycler for phosphorylation and annealing (37□ for 30 min; 95□ for 5 min; decrease temperature down to 25□ at 5□/min). Annealed oligos were mixed with the linearized vector, T4 DNA Ligase, and 10X T4 DNA ligase buffer and incubated at 16°C overnight for ligation.

### Mammalian cell line culture and lipofection

All cell lines (HEK293T, HeLa, U2OS, IMR90) were cultured in Dulbecco’s Modified Eagle Medium (DMEM; Thermo Fisher) supplemented with 10% fetal bovine serum (FBS, VWR), and 1% Penicillin/Streptomycin (P/S, Gibco). All cells were cultured at 37°C in a 5% CO2 air incubator. Routine checks for mycoplasma contamination were performed using the MycoAlert mycoplasma detection kit (Lonza). Lipofection was performed using Lipofectamine 3000 (Thermo Fisher Scientific) according to the manufacturer’s instructions. 50,000 cells per well were seeded in 24-well plates 24 hours prior to lipofection. Cells were transfected with 500 ng Cas nuclease expression plasmid, 150 ng sgRNA expression plasmid, and 1.5 pmol ssODN templates (IDT, Table S2) per well.

### Flow cytometry analysis

Cells were resuspended in FACS buffer (1% BSA in PBS) and analyzed by flow cytometry for BFP-/GFP- cells (end-joining pathways) and BFP-/GFP+ cells (homology-directed repair pathway) 7 days post-transfection (dpt). Flow cytometry was performed on an Attune NxT flow cytometer with a 96-well autosampler (Thermo Fisher Scientific). Data analysis was performed using the FlowJo v10 software.

### Illumina deep sequencing analysis

DNA was extracted 3 dpt using QuickExtract DNA Extraction Solution (Lucigen) and heated at 65□ for 20 min followed by 95□ for 20 min. DNA samples were then amplified with PrimeSTAR GXL DNA Polymerase (Takara Bio) with PCR forward/reverse primers containing Illumina adapter sequences (Table S3) for 30 cycles (98□ for 10 s, 55□ for 15 s, and 68□ for 1 min).

The resulting amplicons were cleaned by adding 25 uL of amplicon to 45 uL of magnetic beads (UC Berkeley Sequencing Core). The samples were placed on a 96-well magnetic plate for 5 min, and the supernatant was removed. The samples were washed twice with 200 uL of 70% ethanol and eluted in 40 uL of Tris-EDTA Buffer (Corning).

The purified samples were sequenced on an Illumina iSeq by QB3 Genomics at UC Berkeley. NGS sequencing reads were analyzed for HDR-mediated modifications and indels using CRISPResso2 (https://crispresso.pinellolab.partners.org) in batch mode using default parameters.

### Western blotting

Cells were resuspended in lysis buffer (20mM Tris pH 7.5, 1mM MgCl2, 1mM CaCl2, 137mM NaCl, 10% Glycerol, 1% NP-40) containing Benzonase (12.5 u/mL) and Protease Inhibitor Cocktail (Roche) and incubated at 4□ for 1 h with rotation. Proteins from whole cell lysates were separated by 4–20% TGX gel (Bio-Rad) and transferred to a nitrocellulose membrane (0.2μm pore size) (Invitrogen). Membranes were blocked and incubated with primary antibodies at 4°C overnight followed by secondary antibodies at room temperature for 1 h. Primary antibodies used for Western blot analysis were anti-FLAG M2 (Sigma, Cat# F3165, RRID:AB_259529) and anti-GAPDH (Cell Signaling Technology, Cat# 2118, RRID:AB_561053). Secondary antibodies were anti-mouse Alexa Fluor™ 647 (Thermo Fisher, Cat# A-31571, RRID:AB_162542) and anti-rabbit Alexa Fluor™ 488 (Thermo Fisher, Cat# A-21206, RRID:AB_2535792). Imaging was performed using the Odyssey imaging system (LI-COR). Double bands were caused by incomplete cleavage at the self-cleaving P2A peptide, leading to an uncleaved byproduct (57).

### ChIP-qPCR

Cells were crosslinked for 5 min in 1% formaldehyde, and the reaction was quenched by the addition of glycine to 125 mM and incubation for 5 min. Cells were washed twice with lysis buffer 1 (50mM HEPES-KOH pH 7.5, 140mM NaCl, 1mM EDTA, 10% glycerol, 0.5% NP-40, 0.25% Triton X-100, 1x protease inhibitor) and lysis buffer 2 (10mM Tris-HCl pH 8.0, 200mM NaCl, 1mM EDTA, 0.5mM EGTA, 1x protease inhibitors), respectively. Then, cells were resuspended in lysis buffer 3 (10mM Tris-HCl pH 8.0, 100mM NaCl, 1mM EDTA, 0.5mM EGTA, 0.1% Na-Deoxycholate, 0.5% N-lauroylsarcosine, 1x protease inhibitors) and sheared by sonication (peak power 140, duty factor 5%, 200 cycles per burst, 900 s treatment time) using a Covaris S2 (UC Berkeley Functional Genomics Laboratory). 500 ug lysate was incubated at 4□ overnight with 2 ug of appropriate antibodies (anti-H3K4me3 (Abcam, Cat# ab8580, RRID: AB_306649), anti-H3K36me3 (Abcam, Cat# ab9050, RRID: AB_306966), anti-H3K36me2 (Abcam, Cat# ab9049, RRID: AB_1280939), and anti-IgG (Abcam, Cat# ab46540, RRID: AB_2614925)) pre-bound to 30 uL of Protein G Magnetic Beads (Thermo Fisher). The IP/bead mixture was washed 5 times with RIPA wash buffer (50mM HEPES-KOH pH 7.5, 500mM LiCl, 1mM EDTA, 1% NP-40, 0.7% Na-Deoxycholate) and eluted from the beads for 30 min at 65 □. Both input and IP samples were reverse crosslinked by incubating at 65□ overnight with shaking. Then, samples were incubated at 37□ for 2 h with RNase A (0.2 mg/mL) and at 55□ for 2 h with Proteinase K (0.2 mg/mL). DNA samples were purified using 400 uL UltraPure™ Phenol:Chloroform:Isoamyl Alcohol (25:24:1) (Thermo Fisher).

Quantitative PCR was performed using QuantStudio 3 real-time PCR system (Thermo Fisher). Reactions were prepared in triplicates containing 2.5 uL of DNA sample, 0.1 uL of 10 mM forward/reverse primers (Table S4), 5 uL of Power SYBR Green Master Mix (Thermo Fisher). qPCR settings were: 50°C for 2 min, 95°C for 2 min, followed by 40 cycles of 95°C for 1 min and 60°C for 30 sec. The % of input DNA was calculated as ((Ct[Input] - log2(Input Dilution Factor)) − Ct[ChIP]) × 100%.

### RNA-seq

We quantified the expression of relevant genes in parental HEK293T cells using RNA-seq. Briefly, total RNA was extracted with TRIzol Reagent (Thermo Fisher Scientific) for one replicate. Strand-specific cDNA was prepared from polyA-enriched mRNA, then 150bp paired-end sequenced on an Illumina NovaSeq platform at Novogene (Novogene Corporation Inc., Sacramento, CA) to a depth of 30 million reads. Transcripts-per-million for each gene were determined with Kallisto (version 0.48.0), using the GRCh38 reference transcriptome and a kmer size of 31.

## Author Contributions

This project was conceived by E.L.-S. and J.A.D. E.L.-S. and E.C. initiated the study and designed and conducted all of the experiments. M.T. performed all bioinformatics analyses. M.S.D. aided in molecular biology experiments. D.C. performed the RNA-sequencing experiments. E.C., E.L.-S., M.T., and J.A.D. wrote the manuscript. All authors reviewed and commented on the manuscript.

## Data availability

All data needed to evaluate the conclusions in the paper are present in the paper and/or the Supporting Information. All ChIP-seq datasets used during this study are publicly available. The RNA-seq dataset used in this study is available under the accession number PRJNA866532 (https://www.ncbi.nlm.nih.gov/sra/PRJNA866532).

