## Supplemental file for "Decorating chromatin for enhanced genome editing using CRISPR-Cas9"

A)

| | PTM | Biological Process | Presumed Role in DNA Repair |
|---|---|---|---|
| PRDM9 | H3K4me3<br>H3K36me3 | Localization of meiotic recombination hotspots | H3K36me3 and H3K4me3 recruit ZCWPW1 to facilitate repair of SPO11-induced DSBs via HR |
| SETD2 | H3K36me3 | DNA repair by HR, mismatch repair, V(D)J recombination, transcription fidelity | H3K36me3 recruits CtIP to promote DSB resection and RPA and RAD51 foci formation |
| SETMAR | H3K4me3<br>H3K36me2 | DNA repair by NHEJ, supression of chromosomal translocations | H3K36me2 stabilizes NSB1 and Ku70 at DSB sites to promote NHEJ repair. |

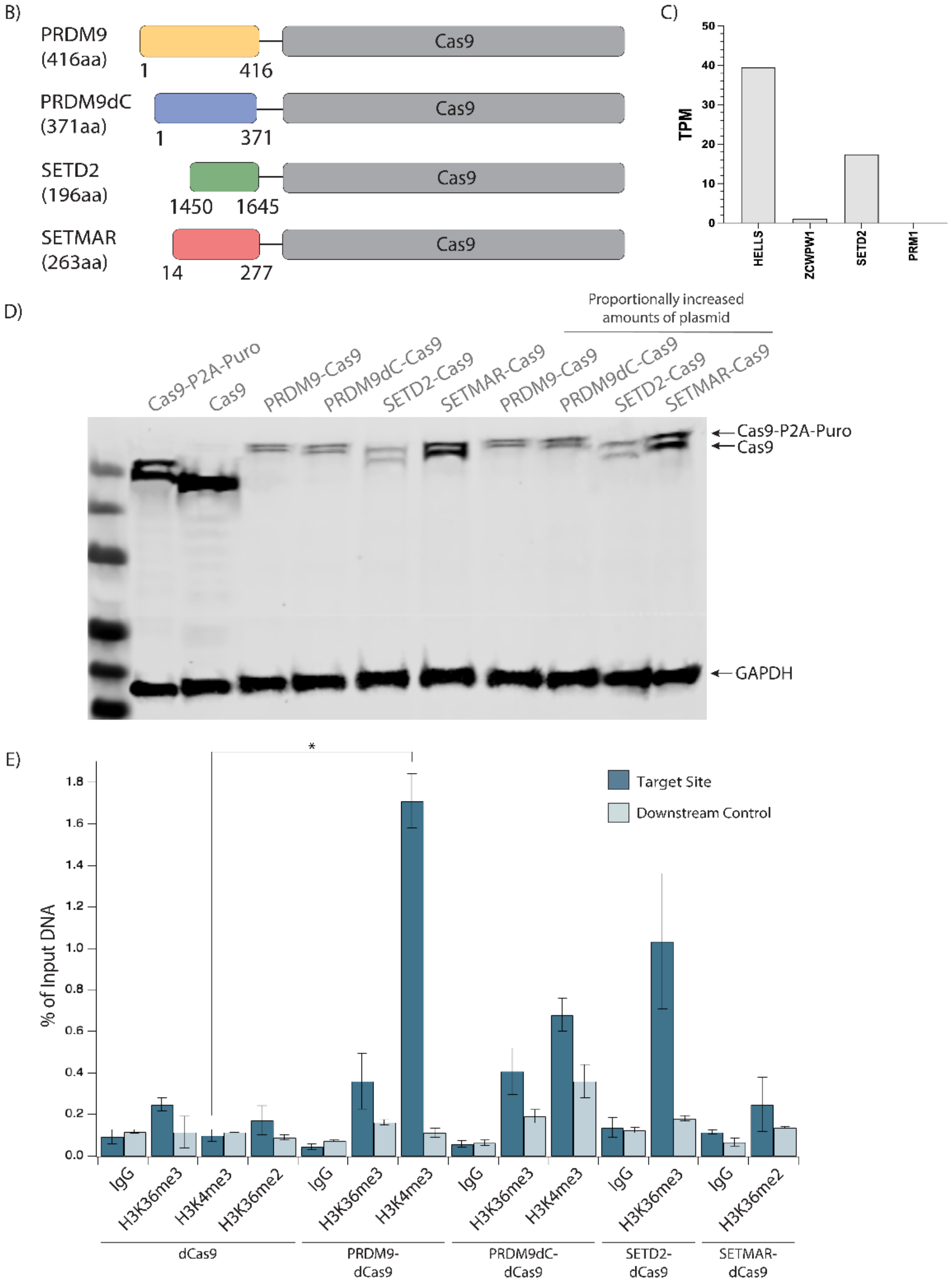

**Fig. S1.** Cas9-methyltransferase fusion protein design to modulate the choice of DNA repair pathway

A) Summary of histone methyltransferases used for Cas9 fusion protein engineering, histone posttranslational modifications (PTM) deposited, biological processes, and their presumed role in DNA repair.
B) Schematic of four Cas9-methyltransferase fusion proteins designed to decorate chromatin to modulate the choice of DNA repair pathway following Cas9-induced DSBs. The histone methyltransferases are fused to Cas9 at the N-terminus, and the amino acid positions of the fused proteins selected for cloning are indicated.
C) Expression levels of relevant genes (HELLS, ZCWPW1) based on RNA-seq experiments in HEK293T cells. Positive control (SETD2) and negative control (PRM1) genes were included.
D) Western blot analysis of fusion protein expression in HEK293T cells. GAPDH antibody was used as a loading control. Double bands were due to incomplete cleavage at the self-cleaving P2A peptide, leading to an uncleaved byproduct (1).
E) H3K36me3, H3K4me3, and H3K36me2 enrichment shown as a percentage of input DNA measured by ChIP-qPCR at site C9 in HEK293T cells at 3 dpt. Cells were transfected with plasmids expressing the dCas9 fusion proteins and sgRNA. Data represents mean ± s.d. (n = 2). $^*P < 0.05$, determined by student's two-tailed t-test.

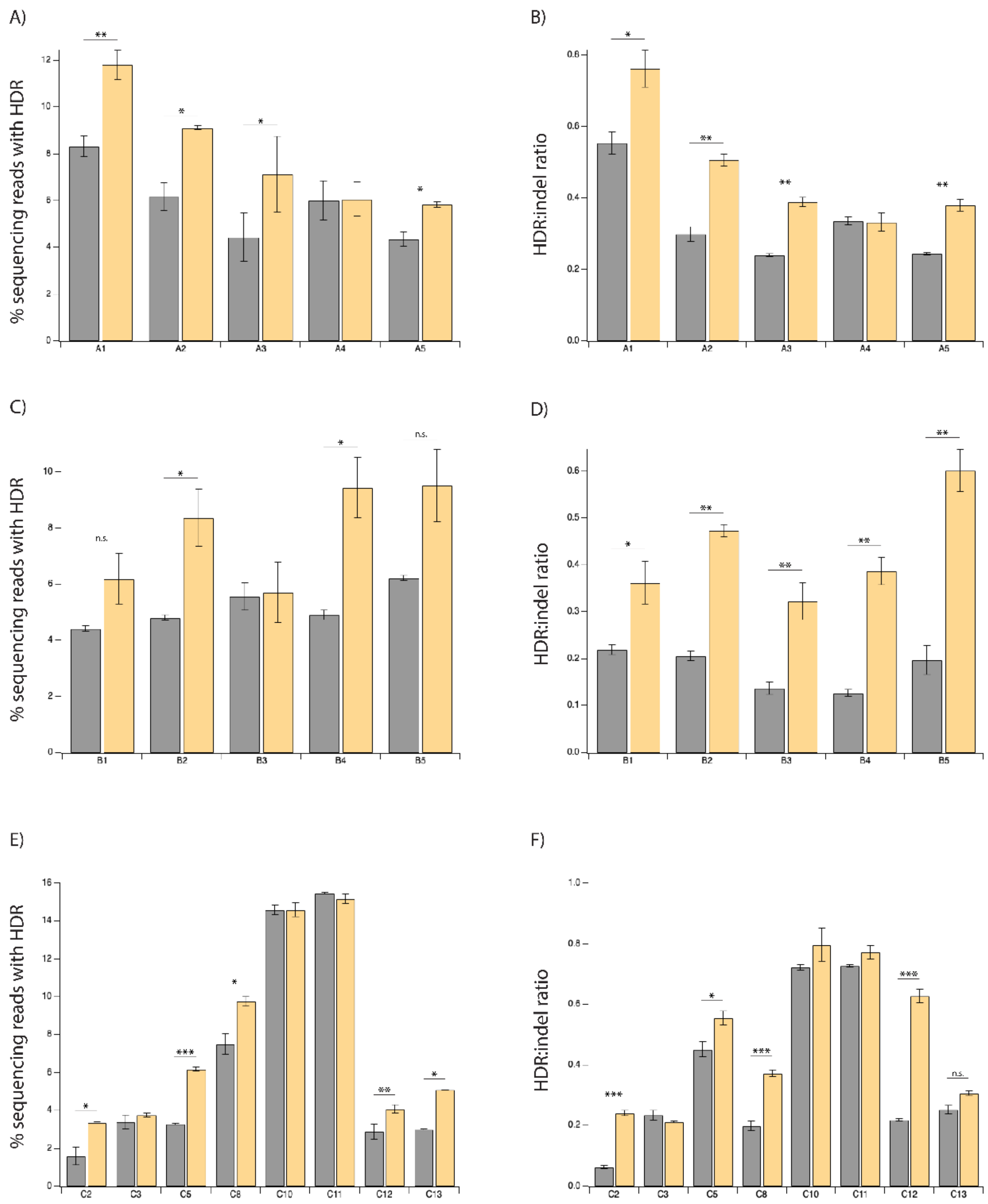


**Fig. S2.** HDR efficiency and HDR:indel ratio improvement by PRDM9-Cas9 at multiple genomic loci

- **A)** HDR frequency and **B)** HDR:indel ratio measured by NGS at five genomic loci enriched with H3K36me3 in HEK293T cells transfected with PRDM9-Cas9 and sgRNA with ssODN template.
- **C)** HDR frequency and **D)** HDR:indel ratio measured by NGS at five genomic loci enriched with H3K36me3 and H3K4me3 in HEK293T cells transfected with PRDM9-Cas9 and sgRNA with ssODN template.
- **E)** HDR frequency and **F)** HDR:indel ratio measured by NGS at eight genomic loci lacking H3K36me3 and H3K4me3 in HEK293T cells transfected with PRDM9-Cas9 and sgRNA with ssODN template.

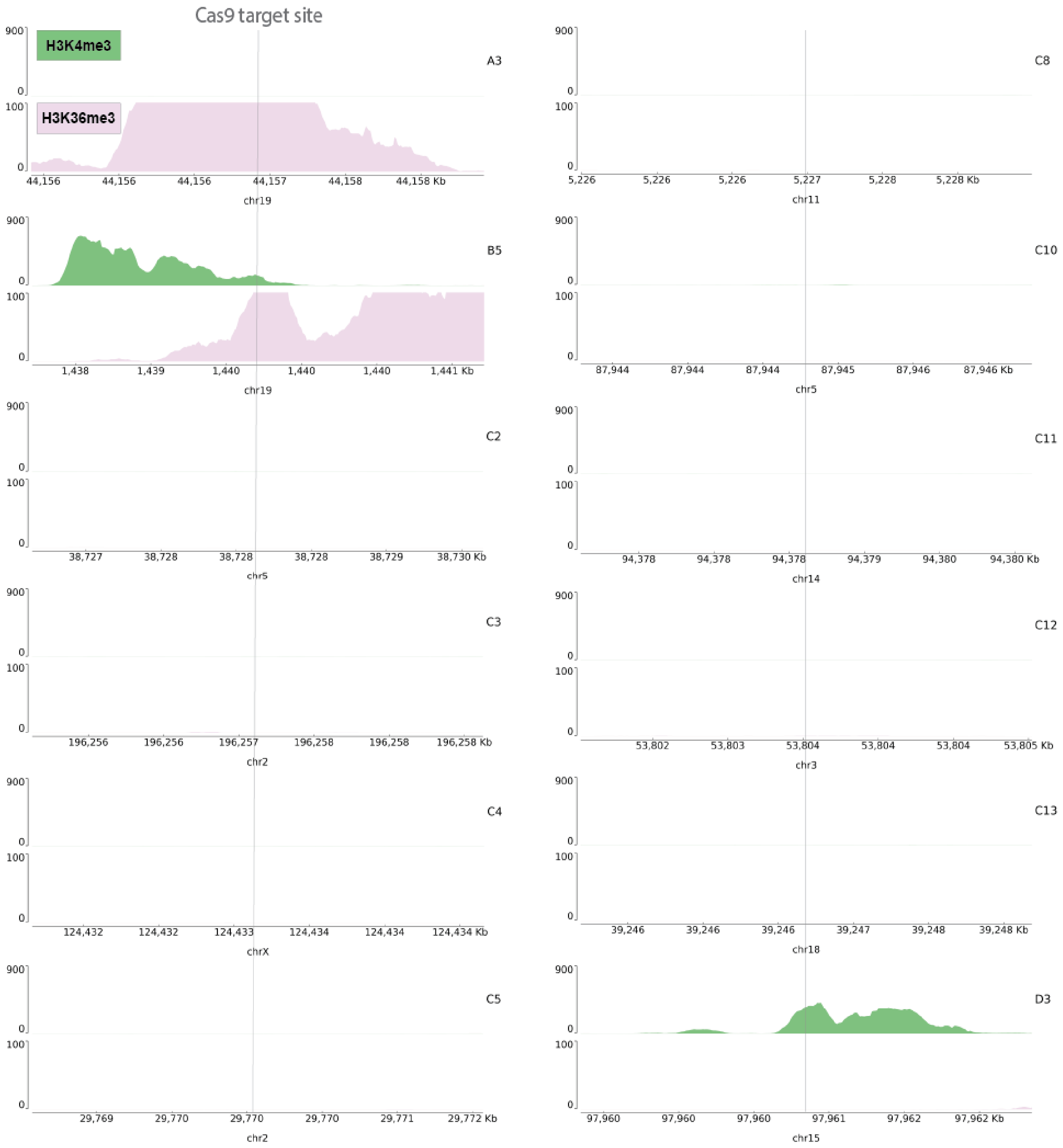


**Fig. S3.** UCSC genome browser tracks showing endogenous levels of H3K4me3 and H3K36me3 enrichment at additional genomic sites

Tracks represent target site ± 1.5 kilobases (kb) in HEK293 cells based on ENCODE ChIP-seq datasets for H3K4me3 (green) and H3K36me3 (pink). Each plot spans 3 kb and the y-axis reports the negative-log p-value for peak enrichment.

A)

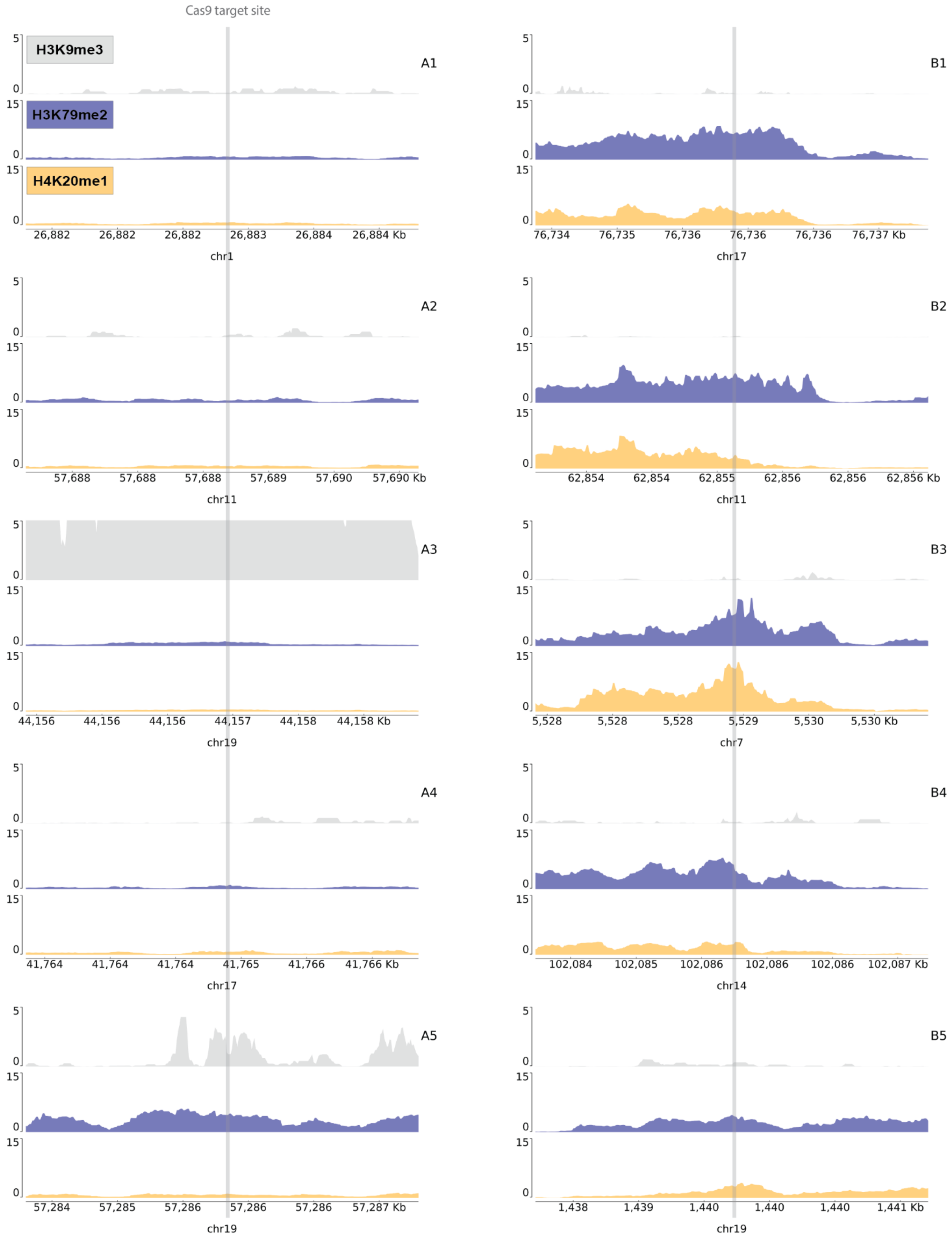

Cas9 target site
H3K9me3
H3K79me2
H4K20me1
A1
B1
26,882 26,882 26,882 26,883 26,884 26,884 Kb
chr1
76,734 76,735 76,736 76,736 76,736 76,737 Kb
chr17
A2
B2
57,688 57,688 57,688 57,689 57,690 57,690 Kb
chr11
62,854 62,854 62,855 62,856 62,856 62,856 Kb
chr11
A3
B3
44,156 44,156 44,156 44,157 44,158 44,158 Kb
chr19
5,528 5,528 5,528 5,529 5,530 5,530 Kb
chr7
A4
B4
41,764 41,764 41,764 41,765 41,766 41,766 Kb
chr17
102,084 102,085 102,086 102,086 102,086 102,087 Kb
chr14
A5
B5
57,284 57,285 57,286 57,286 57,286 57,287 Kb
chr19
1,438 1,439 1,440 1,440 1,440 1,441 Kb
chr19

B)

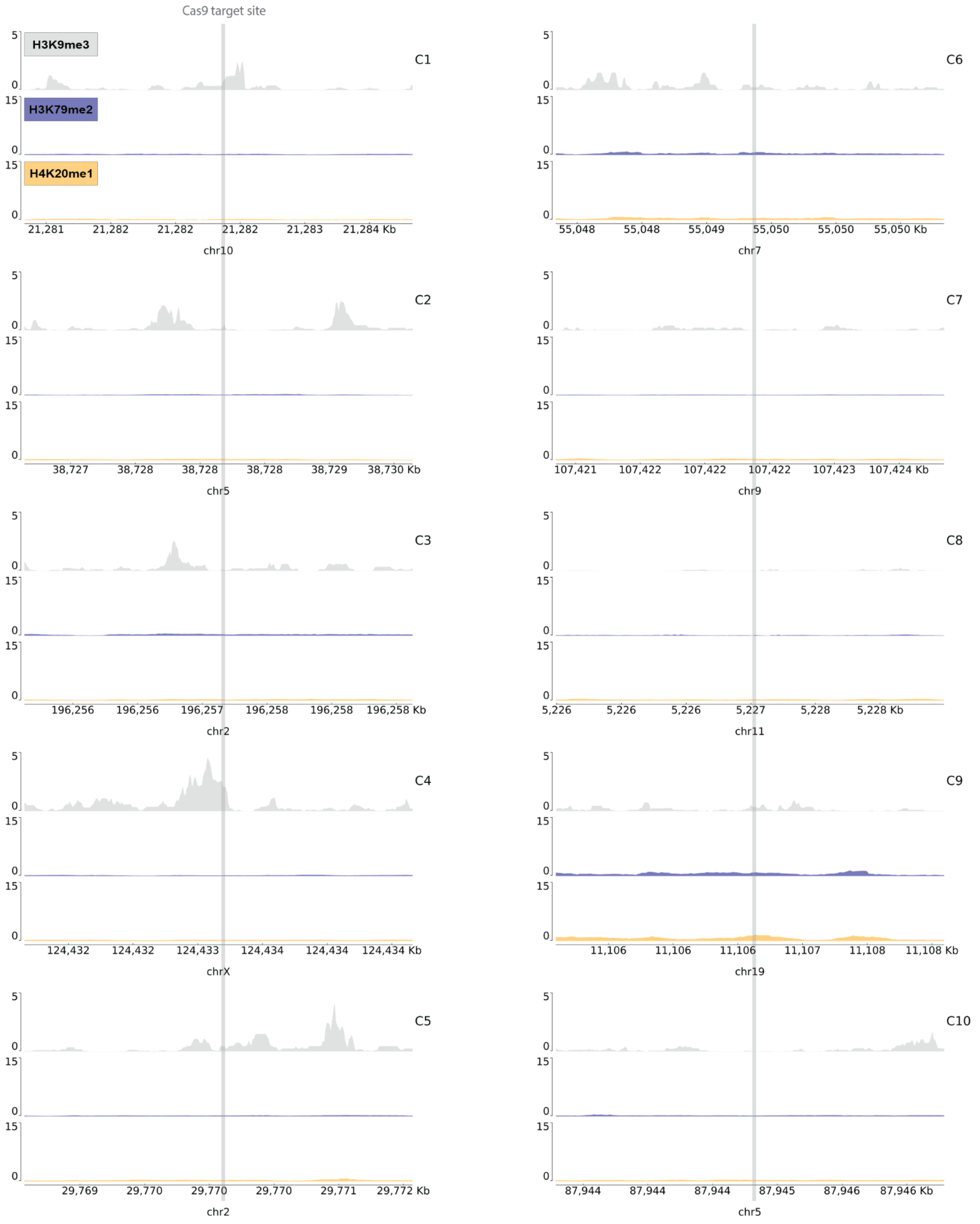
Cas9 target site
H3K9me3
H3K79me2
H4K20me1
C1
21,281 21,282 21,282 21,282 21,283 21,284 Kb
chr10
C6
55,048 55,048 55,049 55,050 55,050 55,050 Kb
chr7
C2
38,727 38,728 38,728 38,728 38,729 38,730 Kb
chr5
C7
107,421 107,422 107,422 107,422 107,423 107,424 Kb
chr9
C3
196,256 196,256 196,257 196,258 196,258 196,258 Kb
chr2
C8
5,226 5,226 5,226 5,227 5,228 5,228 Kb
chr11
C4
124,432 124,432 124,433 124,434 124,434 124,434 Kb
chrX
C9
11,106 11,106 11,106 11,107 11,108 11,108 Kb
chr19
C5
29,769 29,770 29,770 29,770 29,771 29,772 Kb
chr2
C10
87,944 87,944 87,944 87,945 87,946 87,946 Kb
chr5

C)

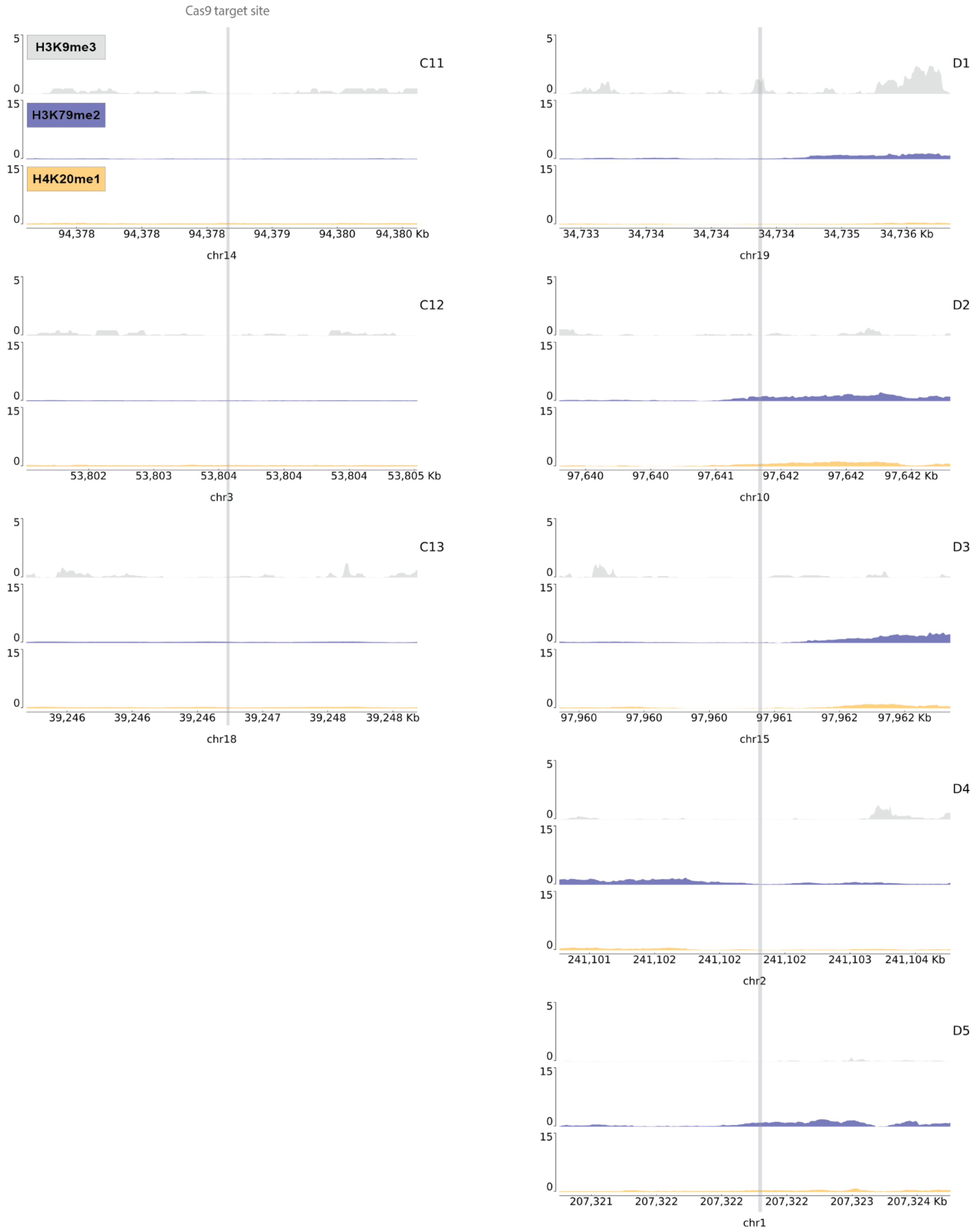
Cas9 target site
H3K9me3
H3K79me2
H4K20me1
C11
94,378 94,378 94,378 94,379 94,380 94,380 Kb
chr14
C12
53,802 53,803 53,804 53,804 53,804 53,805 Kb
chr3
C13
39,246 39,246 39,246 39,247 39,248 39,248 Kb
chr18
D1
34,733 34,734 34,734 34,734 34,735 34,736 Kb
chr19
D2
97,640 97,640 97,641 97,642 97,642 97,642 Kb
chr10
D3
97,960 97,960 97,960 97,961 97,962 97,962 Kb
chr15
D4
241,101 241,102 241,102 241,102 241,103 241,104 Kb
chr2
D5
207,321 207,322 207,322 207,322 207,323 207,324 Kb
chr1

**Fig. S4.** UCSC genome browser tracks showing normalized endogenous levels of H3K9me3, H3K79me2, and H4K20me1 at genomic sites

Tracks represent target site ± 1.5 kilobases (kb) in HEK293 cells based on ENCODE ChIP-seq datasets for H3K9me3 (gray), H3K79me2 (blue), and H4K20me1 (yellow) at **A)** sites A1-A5 and B1-B5, **B)** sites C1-C10, and **C)** sites C11-C13 and D1-D5. Each plot spans 3 kb and the y-axis reports the negative-log p-value for peak enrichment.

**Table S1.** Single guide RNA (sgRNA) sequences and HDR products

| Site | Locus | sgRNA and PAM sequence | Change encoded by ssODN template |
|---|---|---|---|
| A1 | GPN2 | GGACAGCTCTGGATGTCCCAtgg | GGACAGCTCTGGATGTGCCAtgg |
| A2 | ZDHHC5 | GTTGTCACAGACACTGCAGTggg | GTTGTCACAGACACTGGAGTggg |
| A3 | ZNF234 | TATTAAGTGCTGATCTACGAcgg | TATTAAGTGCTGATCTTCGAcgg |
| A4 | JUP | TCAGTCAGCCATGCTCCCCGtgg | TCAGTCAGCCATGCTCGCCGtgg |
| A5 | ZNF460 | GTGGAAGAACTCAATCCTAAggg | GTGGAAGAACTCAATCGTAAggg |
| B1 | SRSF2 | GGAGACCGCAGCTTTAAAGGggg | GGAGACCGCAGCTTTATAGGggg |
| B2 | SNHG1 | CATGGTTTATGCTCTTACAGagg | CATGGTTTATGCTCTTTCAGagg |
| B3 | ACTB | GCTCGTGTGACAAGGCCATGagg | GCTCGTGTGACAAGGCGATGagg |
| B4 | HSP90AA1 | TTCACTGGTTAAGTGAGCGGtgg | TTCACTGGTTAAGTGACCGGtgg |
| B5 | RPS15 | AAAGGCCTCTAAAAACGCAGggg | AAAGGCCTCTAAAAACCCAGggg |
| C1 | NEBL | ACAATGCAAGAGGCTCCGTAggg | ACAATGCAAGAGGCTCGGTAggg |
| C2 | OSMR-DT | ACTGAATTCCTTGAAGACCGggg | ACTGAATTCCTTGAAGTCCGggg |
| C3 | HECW2 | GGATGAGCGTCCCAACCACTggg | GGATGAGCGTCCCAACGACTggg |
| C4 | TENM1 | CTTGGTTGAATCTAATCACGggg | CTTGGTTGAATCTAATGACGggg |
| C5 | ALK | GTCTCTTGCTGGATATGGGAagg | GTCTCTTGCTGGATATCGGAagg |
| C6 | EGFR | GTAATGAAATGAGTTGGGGCagg | GTAATGAAATGAGTTGCGGCagg |
| C7 | | GGCCCAGACTGAGCACGTGAtgg | GGCCCAGACTGAGCAAGTGAtgg |
| | | | GGCCCAGACTGAGCAAGTGATTT (PAM mutation) |
| C8 | HBB | GTAACGGCAGACTTCTCCTCagg | GTAACGGCAGACTTCTCCACagg |
| C9 | LDLR | CAGAGCACTGGAATTCGTCAggg | CAGAGCACTGGAATTAGTCAggg |
| | | | CAGAGCACTGGAATTAGTCAGTG (PAM mutation) |
| C10 | | GAACACAAAGCATAGACTGCggg | GAACACAAAGCAGAGACTGCggg |
| | | | GAACACAAAGCAGAGACTGCGAC (PAM mutation) |
| C11 | SERPINA1 | TGCTGACCATCGACGAGAAAggg | TGCTGACCATCGACAAGAAAggg |
| C12 | CACNA1D | GGAGCAGGAGTATTTCAGTAGTGagg | GGAGCAGGAGTATTTCAGTTGTGagg |
| C13 | | AAGATGCAAGGTTTGTGTCTagg | AAGATGCAAGGTTTGTCTCTagg |
| D1 | ZNF181 | TTGCCCGCTCACCAGCGACGcgg | TTGCCCGCTCACCAGCCACGcgg |
| D2 | PI4K2A | GCATCCCTAAATCCTGCCAGggg | GCATCCCTAAATCCTGGCAGggg |
| D3 | ARRDC4 | GGCAGGAAAGAGTCGCCCGGcgg | GGCAGGAAAGAGTCGCGCGGcgg |
| D4 | MTERF4 | GGCCGAACGCAGCCATAGCGcgg | GGCCGAACGCAGCCATTGCGcgg |
| D5 | CD55 | GCAGCATCGTGTGCTCCACAcgg | GCAGCATCGTGTGCTCGACAcgg |
| BFP-GFP | | GCTGAAGCACTGCACGCCATggg | GCTGAAGCACTGCACGCCGTACG |

**Table S2.** Single-stranded oligonucleotide (ssODN) donor templates

| Site | Template Sequence |
|---|---|
| A1 | CACTAACCAAGCTCCAAAAGTCTGGCTGCCTCTTGGACAGCTCTGGATGTGCCATGGACCCCTGA ACACAGCACATCTGCCAGCTCTTGTCCCTTTCATCC |
| A2 | GTCTTACCCCACCCAGGGTGCTTACCTCCACACAGTTGTCACAGACACTGGAGTGGGAACATCGA GGGGGACGGTAAAAGCGGCAGGTGGCACACCATTTC |
| A3 | TATGGTTTCTCTCCTGTGTGGACCATGCAATGATTATTAAGTGCTGATCTTCGACGGAAGTTCTTAC CACATGTATCACATTTGAATGGTTTCTCCCCAGT |
| A4 | CGGCTGGGGTGGGAGCTCTGCCCAGCTGCCAGGCTCAGTCAGCCATGCTCGCCGTGGCTGATG GCACCCCTATCCCTGCCCCAGGTGGGATGCAGGCCCTG |
| A5 | AAAACAAGGAGGCCATAAAGAATGGGAGACATGAGTGGAAGAACTCAATCGTAAGGGAACAGGGC AGCAGAGTGAGGGGTAGACGTGGTGACTCCACGCTT |
| B1 | TCTGCAATACTGGCCAATATTCTTTTATCAAACAGGAGACCGCAGCTTTATAGGGGGAAAATGCAG ACGTTGGATAAAAACAGCAAGAAATAGTCATTTTC |
| B2 | AGCCAGCGTTACAGTAATGTTCCAGGTAGGTGTACATGGTTTATGCTCTTTCAGAGGAGACCTTGT AGATAACCACTCCATGATGAACACAAAATGACAAG |
| B3 | TAAGACAGTGTTGTGGGTGTAGGTACTAACACTGGCTCGTGTGACAAGGCGATGAGGCTGGTGTA AAGCGGCCTTGGAGTGTGTATTAAGTAGGTGCACAG |
| B4 | ACAGTGAGGACAGACACAGGTAGGCACCTGAACATTCACTGGTTAAGTGACCGGTGGAAGGGTG GGGTGCTGCAACCCTTGGACCCCTGGGGATGCGGTGA |
| B5 | CTCAGACAAGGTGCTGCAGCGTACAGCTCGGGCCAAAGGCCTCTAAAAACCCAGGGGAAGGCAG GTAGGGCGCAGTGGCTCCCGCCTGTGATCCCAGCACT |
| C1 | GGAATTCCTTCATTACAAATATTTATTAGTACCTACAATGCAAGAGGCTCGGTAGGGAGCACAAACC TAAGGAAAGGTGCAAACAAACAAAACCAAGCTTT |
| C2 | GAGCTTATGGTAGAGTTAGAGAAAAGAATGCACAACTGAATTCCTTGAAGTCCGGGGCCTCGACTT AAAATGTGATTATAGGGAGTGGAGTGGAAGGGGTC |
| C3 | TATGCTGTTCCTTTTAATTCTCTGGAGGTTACAGGGATGAGCGTCCCAACGACTGGGTAGAGCAAA GCATTATTTTACTATGGCTTGCCTACATTTGAGGG |
| C4 | TTACATCACTGTCATGTTAGGGTTTTCTGTTATTCTTGGTTGAATCTAATGACGGGGAAGACAGTGC TCTGTTAGGCTTTGGTCTTGACTGGAGTTACTCA |
| C5 | ATTTCTCCAATTCCAGATAGTAAAGAAGCCTTTGGTCTCTTGCTGGATATCGGAAGGGGATTTGTGG ATCCAGCTGCTTCTTGAATGAACTTTCATCAAGG |
| C6 | CTTCAGGAGAAATAATGAAGAAAGAGGCGGTTTGGTAATGAAATGAGTTGCGGCAGGAGATTACG GTCATTTCAAGTTATATTCAAATTCTCTTCTTTTTG |
| C7 (lacks PAM mutation) | GCTTCTCCAGCCCTGGCCTGGGTCAATCCTTGGGGCCCAGACTGAGCAAGTGATGGCAGAGGA AAGGAAGCCCTGCTTCCTCCAGAGGGCGTCGCAGGAC |
| C7 (contains PAM mutation) | GCTTCTCCAGCCCTGGCCTGGGTCAATCCTTGGGGCCCAGACTGAGCAAGTGATTTCAGAGGAA AGGAAGCCCTGCTTCCTCCAGAGGGCGTCGCAGGAC |
| C8 | ACTTCATCCACGTTCACCTTGCCCCACAGGGCAGTAACGGCAGACTTCTCCACAGGAGTCAGATG CACCATGGTGTCTGTTTGAGGTTGCTAGTGAACAC |
| C9 (lacks PAM mutation) | ATCAACACACTCTGTCCTGTTTTCCAGCTGTGGCCACCTGTCGCCCTGACTAATTCCAGTGCTCT GATGGAAACTGCATCCATGGCAGCCGGCAGTGTGA |
| C9 (contains PAM mutation) | ATCAACACACTCTGTCCTGTTTTCCAGCTGTGGCCACCTGTCGCACTGACTAATTCCAGTGCTCT GATGGAAACTGCATCCATGGCAGCCGGCAGTGTGA |

| | |
|---|---|
| C10 (lacks PAM mutation) | TTTTCCAGCCCGCTGGCCCTGTAAAGGAAACTGGAACACAAAGCAGAGACTGCGGGGCGGGCC AGCCTGAATAGCTGCAAACAAGTGCAGAATATCTGAT |
| C10 (contains PAM mutation) | TTTTCCAGCCCGCTGGCCCTGTAAAGGAAACTGGAACACAAAGCAGAGACTGCGACGCGGGCCA GCCTGAATAGCTGCAAACAAGTGCAGAATATCTGAT |
| C11 | CATGGGTATGGCCTCTAAAAACATGGCCCCAGCAGCTTCAGTCCCTTTCTTGTCGATGGTCAGCA CAGCCTTATGCACGGCCTGGAGGGGAGAGAAGCAG |
| C12 | ATGGCTATTTCAGGGACCCCCACTGCTTGGGGGAGCAGGAGTATTTCAGTTGTGAGGAATGCTAC GAGGATGACAGCTCGCCCACCTGGAGCAGGTGAGCT |
| C13 | AGAACTGGAATCACTATATGTTTATGTGAAAAAAATGAACCTAGAGACAAACCTTGCATCTTTCAAA AAATTAACCAAAAATAGATTGTAGACATAAATG |
| D1 | TCTGGGCAGTGTCGTCCCCAGAGGTCGGCGGCCGTTGCCCGCTCACCAGCCACGCGGGGCGC CTGCGGGGACGGTGAGGCCCTGCTGAGGACTCCGGCCAG |
| D2 | TTCCGCAGAGCGTCCCCAGCTCCAGTCCTCAGTGGCATCCCTAAATCCTGGCAGGGGCTGGTAC TGTGAAGTTTCTCCCTTTTCCCTCTGGCTGTGCGAAG |
| D3 | GCGACCCCGGCTCCGGGCCTCTGCCGACCTCAGGGGCAGGAAAGAGTCGCGCGGCGGGATGG GCGGGGAGGCTGGGTGCGCGGCGGCCGTGGGTGCCGAGG |
| D4 | AGAAGCCCGCGCGCCCAGCTCGAGCTTACCTGACGGCCGAACGCAGCCATTGCGCGGAGAAGA TGGCAGCAGTTACGGCGCCGGAAGCAGCGGTCCTCCCC |
| D5 | AATGCCACACCTGAAAGAGATGACATGCAAGTTTGCAGCATCGTGTGCTCGACACGGCTGGACTC TGTCGAGAGTGGGGAACGGTCAGCGGTCTGGCCTCC |
| BFP-GFP | GCCACCTACGGCAAGCTGACCCTGAAGTTCATCTGCACCACCGGCAAGCTGCCCGTGCCCTGG CCCACCCTCGTGACCACCCTGACGTACGGCGTGCAGTGCTTCAGCCGCTACCCCGACCACATGA |

**Table S3.** PCR primers for constructing next-generation sequencing libraries

| Site | Primer |
|---|---|
| A1 (for) | TCCCTACACGACGCTCTTCCGATCTAGGAGGGAGCAGGTTTGTCA |
| A1 (rev) | GTTCAGACGTGTGCTCTTCCGATCTGGACACCTACCCATGACCTTG |
| A2 (for) | TCCCTACACGACGCTCTTCCGATCTGATGATTTCCGAGCTCCCCT |
| A2 (rev) | GTTCAGACGTGTGCTCTTCCGATCTGGATGAAGACAATGAAGGTTAAGCA |
| A3 (for) | TCCCTACACGACGCTCTTCCGATCTAGGGCCTTCATACATGCTTCC |
| A3 (rev) | GTTCAGACGTGTGCTCTTCCGATCTACACTTACCACAGTCCTCACAT |
| A4 (for) | TCCCTACACGACGCTCTTCCGATCTGCTGGTGAGTATGATGGCCTTG |
| A4 (rev) | GTTCAGACGTGTGCTCTTCCGATCTCTGTTGCTGGTCAGGTGCTT |
| A5 (for) | TCCCTACACGACGCTCTTCCGATCTGCTATTTGGAGTTCCCTCCC |
| A5 (rev) | GTTCAGACGTGTGCTCTTCCGATCTAGGAAATGAAGAACACCCCCG |
| B1 (for) | TCCCTACACGACGCTCTTCCGATCTAGCTTAGATAATAATGGCTGTTTCG |
| B1 (rev) | GTTCAGACGTGTGCTCTTCCGATCTAGAAACTGGTCCCTGGAGGA |
| B2 (for) | TCCCTACACGACGCTCTTCCGATCTGATGTTCAGCCCACAAGAGC |
| B2 (rev) | GTTCAGACGTGTGCTCTTCCGATCTACATCACTTGAAAGTTCAGCCA |
| B3 (for) | TCCCTACACGACGCTCTTCCGATCTTCTGGTGTTTGTCTCTCTGACTA |
| B3 (rev) | GTTCAGACGTGTGCTCTTCCGATCTGGACATGCAGAAAGTGCAAAG |
| B4 (for) | TCCCTACACGACGCTCTTCCGATCTAGTAACTGTGATCACCAAACATAAC |
| B4 (rev) | GTTCAGACGTGTGCTCTTCCGATCTGATCGTTGGGCAAACACAAA |
| B5 (for) | TCCCTACACGACGCTCTTCCGATCTTGAACTCCTGGGCTCAA |
| B5 (rev) | GTTCAGACGTGTGCTCTTCCGATCTCGTGTGACTGCAGAGATTC |
| C1 (for) | TCCCTACACGACGCTCTTCCGATCTAATGGATGAGCTCTCTGCGG |
| C1 (rev) | GTTCAGACGTGTGCTCTTCCGATCTACACTATGGTAGCCGTCCCT |
| C2 (for) | TCCCTACACGACGCTCTTCCGATCTGCTGAGAAGGACTTCAAGAGAT |
| C2 (rev) | GTTCAGACGTGTGCTCTTCCGATCTCCATATGTGAGAAATAAACAGTCCTAC |
| C3 (for) | TCCCTACACGACGCTCTTCCGATCTAGTAGTTTAGAAAGGAAACTGGCAA |
| C3 (rev) | GTTCAGACGTGTGCTCTTCCGATCTAAAACCTTGCCTGAAAGGCTCA |
| C4 (for) | TCCCTACACGACGCTCTTCCGATCTAGGTGATTCATCCAGAGGTGTA |
| C4 (rev) | GTTCAGACGTGTGCTCTTCCGATCTAAACCACACATCTAGCCTGG |
| C5 (for) | TCCCTACACGACGCTCTTCCGATCTGGTAGTTTACCCTCCTCCTCTA |
| C5 (rev) | GTTCAGACGTGTGCTCTTCCGATCTAGGCATTGATTGTGCTATCTCTA |
| C6 (for) | TCCCTACACGACGCTCTTCCGATCTGGTATCTGCCCAGAAAGCTCTA |
| C6 (rev) | GTTCAGACGTGTGCTCTTCCGATCTGCTGAGTGCAGTGGGAGAGA |
| C7 (for) | GCTCTTCCGATCTGGAAACGCCCATGCAATTAGTC |
| C7 (rev) | GCTCTTCCGATCTCTTGTCAACCAGTATCCCGGTG |
| C8 (for) | GCTCTTCCGATCTCTTGATACCAACCTGCC |
| C8 (rev) | GCTCTTCCGATCTTAAAAGTCAGGGCAGAGCCA |

| | |
|---|---|
| C9 (for) | GCTCTTCCGATCTGCCCTGCTTCTTTTTCTCTGGT |
| C9 (rev) | GCTCTTCCGATCTACCATTAACGCAGCCAACTTCA |
| C10 (for) | GCTCTTCCGATCTCCAGCCCCATCTGTCAAACT |
| C10 (rev) | GCTCTTCCGATCTGAATGGATTCCTTGGAAACAATG |
| C11 (for) | GCTCTTCCGATCTTTTGTTGAACTTGACCTCGGGG |
| C11 (rev) | GCTCTTCCGATCTTCAGGGAGGGAGAGGATGTG |
| C12 (for) | TCCCTACACGACGCTCTTCCGATCTAGGAGATGAACAGCTCCCAACTA |
| C12 (rev) | GTTCAGACGTGTGCTCTTCCGATCTGCTTTTTAGGGGAAGCCCAAGT |
| C13 (for) | GCTCTTCCGATCTTGACACACACACACATGAAAT |
| C13 (rev) | GCTCTTCCGATCTCCAAACGCAAGATCATGTAGA |
| D1 (for) | TCCCTACACGACGCTCTTCCGATCTGTCCGCCCCGCTGTAG |
| D1 (rev) | GTTCAGACGTGTGCTCTTCCGATCTACCCAGGGCCTGAGCC |
| D2 (for) | TCCCTACACGACGCTCTTCCGATCTGGTGAGTGCGGGGGTG |
| D2 (rev) | GTTCAGACGTGTGCTCTTCCGATCTTCCTCCCCCACCAAGTAGG |
| D3 (for) | TCCCTACACGACGCTCTTCCGATCTCCTTACCCTGCCGCGAG |
| D3 (rev) | GTTCAGACGTGTGCTCTTCCGATCTCTCGTCCTCGAACACCAGAC |
| D4 (for) | TCCCTACACGACGCTCTTCCGATCTGGGCCGGAAGTGAAGGAC |
| D4 (rev) | GTTCAGACGTGTGCTCTTCCGATCTACCTGGCCGTGAAACACTC |
| D5 (for) | TCCCTACACGACGCTCTTCCGATCTAGGGAGGGCTCAAAGAGACT |
| D5 (rev) | GTTCAGACGTGTGCTCTTCCGATCTTTCAAGACACAAGCCCCCTT |

**Table S4.** ChIP-qPCR primers

| Site | Primer |
|---|---|
| C7 (for) | CGCCTGTGATGGGCTAATTG |
| C7 (rev) | GGCCTCTCCAGCCTCATTTG |
| C7 control (for) | CCACATTGGCTCCACCCATC |
| C7 control (rev) | CTTGGTTGTGTCACGTGGTT |
| C9 (for) | GTTGGCTGCGTTAATGGTGA |
| C9 (rev) | TCATTTGCAAGCAGCAAGGC |
| C9 control (for) | AGGCTGAGACAGGAGAATCA |
| C9 control (rev) | GAATGCAATGGCACGATATTGG |
